# Opposing plasticity mechanisms enhance visual inputs that violate their context

**DOI:** 10.1101/2024.06.10.598181

**Authors:** Koen Seignette, Leander de Kraker, Matúš Halák, Paolo Papale, Owen Dorlandt, Lucy S. Petro, Jorrit S. Montijn, Matthew W. Self, Katharina A. Wilmes, Matthew E. Larkum, Pieter R. Roelfsema, Lars F. Muckli, Christiaan N. Levelt

**Affiliations:** Department of Molecular Visual Plasticity, Netherlands Institute for Neuroscience, Amsterdam, the Netherlands; Department of Biology, Humboldt Universität zu Berlin, Berlin, Germany; Department of Vision & Cognition, Netherlands Institute for Neuroscience, Amsterdam,the Netherlands; Department of Medicine and Surgery, Neuroscience Unit, University of Parma, Parma, Italy; Centre for Cognitive Neuroimaging, School of Psychology and Neuroscience, College of Medical, Veterinary and Life Sciences, University of Glasgow, Glasgow, UK; Imaging Centre for Excellence (ICE), College of Medical, Veterinary and Life Sciences, University of Glasgow, UK; Department of Cortical Structure & Function, Netherlands Institute for Neuroscience, Amsterdam, the Netherlands; College of Medical, Veterinary and Life Sciences, University of Glasgow, UK; Institute of Neuroinformatics, ETH Zürich and University of Zürich, Zürich, Switzerland; Laboratory of Visual Brain Therapy, Sorbonne Université, Institut National de la Santé et de la Recherche Médicale, Centre National de la Recherche Scientifique, Institut de la Vision, Paris, France; Department of Integrative Neurophysiology, Centre for Neurogenomics and Cognitive Research, VU University, Amsterdam, the Netherlands; Department of Neurosurgery, Academic University Medical Center, University of Amsterdam, Amsterdam, the Netherlands; Department of Molecular and Cellular Neurobiology, Center for Neurogenomics and Cognitive Research, VU University Amsterdam, Amsterdam, the Netherlands

**Keywords:** Visual cortex, predictive processing, top-down, feedback, plasticity, saliency, learning, occlusion, prediction error, two-photon calcium imaging, computational model, context contrasting

## Abstract

To navigate complex environments, the mammalian visual system prioritizes informative inputs by comparing incoming sensory signals with learned contextual information. While this process is often conceptualized as a difference computation, cortical circuitry poses fundamental challenges: visual contextual feedback is predominantly amplifying, and whether inhibitory circuits support selective subtraction is unknown. Here, we show how layer 2/3 pyramidal neurons in mouse primary visual cortex can implement this comparison through plasticity that acts in opposite directions on responses to direct visual stimulation and contextual signals.

To dissociate these components, we presented full images to mice or images where one quadrant was occluded, removing direct visual stimulation while preserving surrounding contextual information, and measured pyramidal cell responses to these stimuli before and after visual familiarization.

After familiarization to non-occluded images, responses to direct visual inputs were selectively reduced, whereas contextual responses to occluded images strengthened. This strengthening generalized across natural scenes, allowing contextual inputs to also drive responses to novel occluded images and amplify responses to novel non-occluded images. A computational model reproduced these changes by weakening feedforward inputs and strengthening both feedforward-driven inhibition and contextual feedback in response to repeated pyramidal cell activity.

This context-contrasting mechanism produced a conserved structure of responses to occluded and non-occluded images across mice and previously acquired monkey and human data. Together, these results suggest a mechanism for saliency assignment that exploits the compartmental organization of cortical pyramidal neurons: they learn the generalized statistical structure of familiar contexts, allowing selective amplification of improbable inputs or omissions.

## INTRODUCTION

Visual scenes contain far more information than can be processed rapidly. The visual system must therefore prioritize the most informative elements – features that deviate from their surroundings and learned contextual expectations^1–4^. How cortical circuits implement such comparisons remains a central challenge in neuroscience research^5–7^. Theoretical accounts often propose that deviations are detected through subtractive or difference-like computations between feedforward input and contextual feedback^8–13^. This mechanism may indeed account for attenuation of self-motion related responses^14^. However, how such selective subtraction could be implemented by cortical circuits remains unclear. Visual feedback is predominantly amplifying, and although feedback can recruit inhibitory circuits^15^, it is not known whether these provide the feature-specific inhibition required to subtract complex contextual inputs from incoming sensory signals. This raises a fundamental question: how does the cortex compare feedforward visual input with complex contextual signals within the constraints of cortical circuitry?

Addressing this question requires experimentally dissociating direct sensory drive from contextual influences. In primary visual cortex (V1), neurons respond not only to stimuli within their classical receptive fields (cRFs), but also to contextual signals conveyed by recurrent and feedback connections^16^, giving rise to extraclassical receptive field (ecRF) responses^17–20^. Partially occluded visual stimuli provide a powerful approach to disentangle these inputs as they eliminate feedforward input to a retinotopic region of V1 while preserving surrounding visual context^21^. Human fMRI studies^22–26^ and electrophysiological recordings in non-human primates^27^ have shown that occluded regions can evoke robust, image-specific responses in V1, driven by information from beyond the classical receptive field. These responses reflect contextual inputs rather than modulation of direct sensory input, making occlusion paradigms well suited for isolating contextual signals and studying their experience-dependent modulation.

Here, we used two-photon calcium imaging to record the activity of layer 2/3 pyramidal cells (PyCs) in mouse V1 during presentation of non-occluded and partially occluded natural images, before and after visual familiarization. This approach allowed us to ask whether experience alters direct and contextual inputs differently at the level of individual neurons, and whether such changes can account for experience-dependent assignment of saliency to sensory inputs. Our results support a mechanism for saliency assignment that aligns with cortical circuitry, in which PyC connectivity is dynamically optimized to emphasize contrast between feedforward and contextual inputs.

## RESULTS

### Studying the impact of visual experience on feedforward and contextual responses

To assess neural responses to non-occluded and occluded scenes, we recorded calcium signals from L2/3 PyCs in V1 of awake mice using wide-field (WF) and two-photon (2P) imaging while presenting six different natural images in both conditions (Fig. 1A–C). The luminance of the occluded area was equal to that of the gray screen during the intertrial interval, ensuring that neurons with cRFs within the occluded area did not experience any change in feedforward drive when the contextual stimuli were presented (Fig. 1A). Mice were naive to visual input other than normal visual experience within the animal facility.

**Figure 1.**
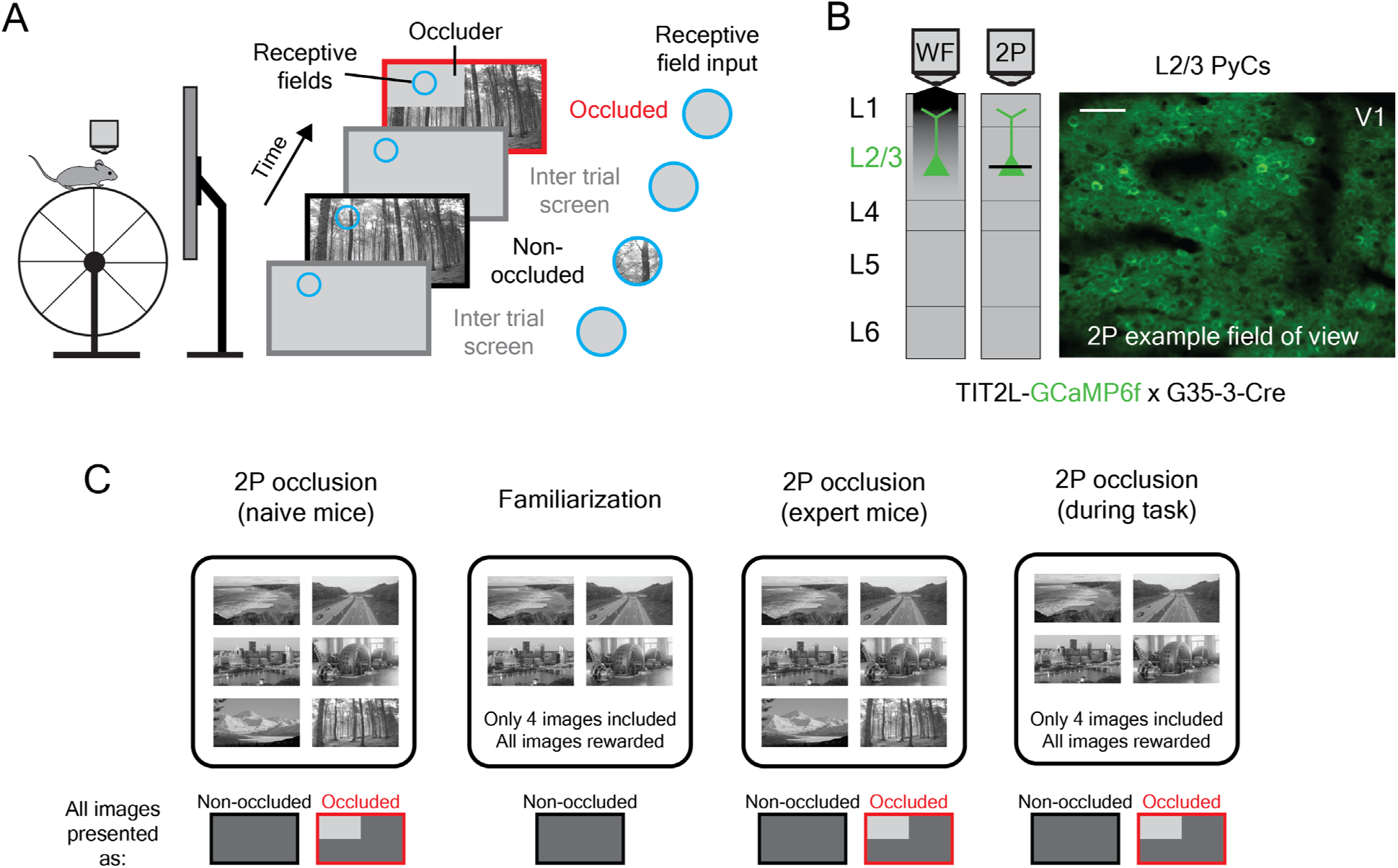
Experimental setup to study plasticity of visual and contextual responses across cell types in mouse V1. A) Schematic of experimental setup. Mice were head-fixed on a running wheel centered in front of a screen displaying non-occluded and occluded natural scenes. B) Imaging approach showing wide-field (WF) and two-photon (2P) calcium imaging in L2/3 PyCs in V1 of TIT2L-GCaMP6f x G35-3 mice. Scale bar, 100 µm. C) Schematic of calcium imaging and behavioral training steps. ‘Naive’ mice were subjected to a WF population RF mapping session followed by retinotopically targeted WF and 2P occlusion experiments. 2P sessions were performed before (naive) and after (expert) a behavioral training phase to familiarize mice with the natural scenes. For behavioral training we used a subset (4/6) of the images, without occlusion. One additional 2P imaging session with familiar images was performed while the mice were engaged in the task (during task).

To test the effect of visual experience on these responses, we familiarized the mice to a subset of the images in a simple stimulus detection task, allowing us to compare neural responses in naive and experienced animals. During this familiarization process, which lasted 2-4 weeks, we presented only four out of the six images to the mice, and only their non-occluded versions. To ensure engagement, mice received a small water reward if they licked within the time window 1–2 s after disappearance of any image. Mice learned to associate the stimuli with a reward, leading to anticipatory-licking and high hit rates (Fig. S1A-B). After the familiarization procedure, we again assessed visual responses to the non-occluded images.

During these passive imaging sessions before (“naive mice”) and after (“expert mice”) familiarization, mice were not performing the task but passively viewed the stimuli and no reward was given. Excluding two out of six images during familiarization allowed us to compare experience-dependent changes in neuronal responses to “familiar” (i.e. used for training) and “novel” (i.e. not used for training) images. To test the influence of engagement, we also recorded responses in one extra 2P session during task performance (“during task”). During this session, occluded and non-occluded images were shown, but the “novel” images were not included.

We performed calcium imaging in L2/3 PyCs by recording in TIT2L-GCaMP6f x G35-3-Cre mice, in which Cre is expressed in cortical excitatory neurons^28^ (Fig. 1B). We targeted our recordings to the region of V1 that retinotopically represented the occluded quadrant of the natural scenes. To achieve this, we first used WF calcium imaging to perform a population receptive field (pRF) mapping^29^ session immediately followed by the occlusion experiment (S1C).

### Experience and engagement drive plasticity of responses to non-occluded and occluded images in opposite directions

We used 2P calcium imaging to record the responses of individual L2/3 PyCs. We studied how these responses depended on visual experience by recording the activity of neurons before and after familiarization with the non-occluded versions of the natural scenes, and during task performance.

Using the WF pRF map, we targeted our 2P imaging to the region of V1 representing the occluder (Fig. 2A). We then mapped the cRF locations of all recorded neurons using sparse noise stimuli and applied strict criteria to only include neurons in all subsequent analyses with high quality cRFs within the occluded region (see Materials and Methods and Figs. 2B and S2). Within this population, both non-occluded and occluded scenes elicited robust responses (Figs. 2A–E). Thus, as previously observed in humans^23,25^ and monkeys^27^, a subset of PyCs in mouse V1 respond to the occluded natural scenes while receiving no feedforward input from the dorsal lateral geniculate nucleus (dLGN), demonstrating that these responses are driven by visual context.

**Figure 2.**
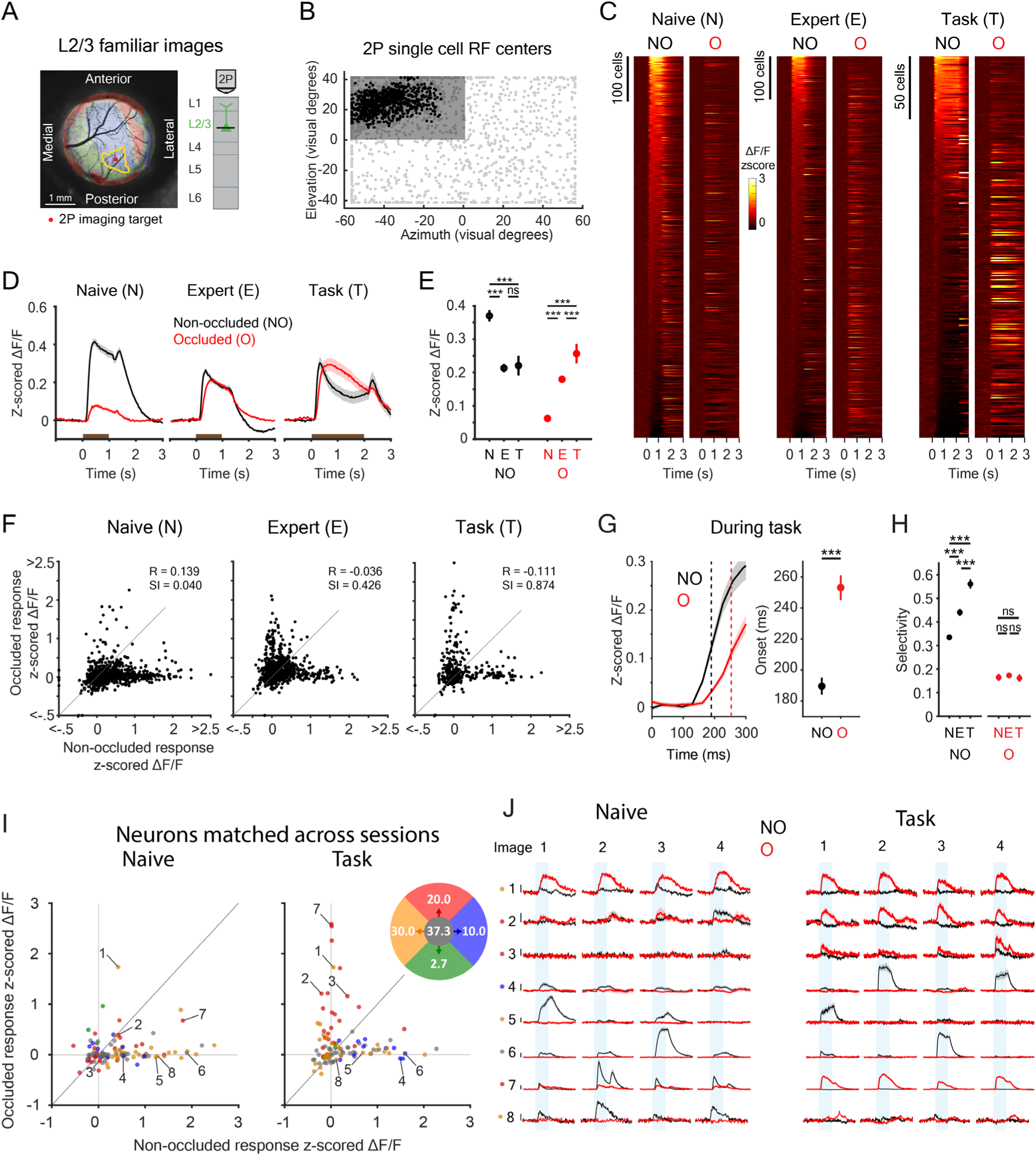
Experience shapes L2/3 responses to familiar feedforward and contextual inputs. A) WF field-sign map showing V1 (blue) and higher visual areas (red) as obtained by population receptive field mapping for an example mouse (same as Figure 1D). Green areas represent regions for which the sign could not be determined. The yellow outline indicates the area in V1 that represents the occluded quadrant on the screen. 2P imaging was targeted in the center of this region (red dot). B) Visual field representation of the stimulus screen showing single cell RF centers. Gray dots are excluded RFs and black dots included RFs. C) Heatmap of z-scored ΔF/F responses for individual neurons across conditions and stimuli. Neurons are sorted within each condition (Naive/Expert/Task) based on their average response (t = 0.2-1 s) to the non-occluded stimuli. N = 841 neurons (Naive), 884 neurons (Expert) and 300 neurons (Task) from 6 mice. NO: non-occluded, O: occluded. D) Z-scored ΔF/F traces averaged across neurons and stimuli for all conditions. Traces represent mean ± SEM over neurons. Brown bars indicate stimulus epoch. E) Mean response strength across stimuli and conditions (mean ± SEM, averaged between t = 0.2-1 s). LMEM with post hoc Tukey HSD for all comparisons, ***: p<0.001, ns: not significant. F) Scatter plot showing average z-scored ΔF/F responses (t = 0.2-1 s) to the non-occluded (x-axis) and occluded (y-axis) stimulus across conditions. N = 841 neurons (Naive), 884 neurons (Expert) and 300 neurons (Task) from 6 mice. R, correlation coefficient; SI, separation index. G) Left: zoom of mean response to NO and O images across neurons during task performance (mean ± SEM). Dotted lines represent mean onset times. Right: quantification of onset times (NO: n = 142 neurons, O: n = 174 neurons). LMEM for all comparisons, ***: p<0.001. H) Stimulus selectivity (lifetime sparseness) for responses to NO and O stimuli. NO: N = 382 neurons (Naive), 223 neurons (Expert) and 79 neurons (Task) from 6 mice. O: N = 79 neurons (Naive), 182 neurons (Expert) and 79 neurons (Task) from 6 mice. LMEM with post hoc Tukey HSD for all comparisons, ***: p<0.001, ns: not significant. I) Scatter plots showing average z-scored ΔF/F responses (t = 0.2-1 s) to the non-occluded (x-axis) and occluded (y-axis) stimulus of chronically matched neurons in naive mice and mice performing the task, color coded for the dominant direction in which their responses transitioned between the naive and the task condition (yellow=reduced NO, blue=increased NO, green=decreased O, red=increased O response, gray=stable). N = 110 neurons from 6 mice. Numbers of the indicated neurons correspond to the neurons in J. The colored disc displays the percentages of the neurons transitioning in each direction. J) Neuronal responses of individual, chronically matched example neurons in naive mice (left) and mice performing the task (right). NO, non-occluded; O, occluded. Scale bar,z 1 z-scored ΔF/F. Traces represent mean ± SEM over trials. The shaded bars indicate stimulus epochs.

We first examined PyC responses to familiar images repeatedly presented during training (Fig. 2C–E). Statistical comparisons were performed using a linear mixed-effects model (LMEM), which accounts for dependencies in the data and variability across mice. In naive mice, a subset of neurons responded reliably to occluded stimuli (4% with z-scored responses > 0.5; Fig. 2C), whereas responses to non-occluded stimuli were more widespread (30% responsive neurons).

Although population-averaged responses measured with two-photon calcium imaging were relatively small (Fig. 2D), this primarily reflected the relatively small fractions of neurons responding to each image rather than weak responses in individual responsive neurons. Among responsive neurons (>0.5 z-scored response), mean response amplitudes were 0.95 z-score (ΔF/F ≈ 9%) for non-occluded images and 0.97 z-score (ΔF/F ≈ 9.5%) for occluded images (Fig. 2C, F), comparable to natural-image responses reported previously

Visual experience had profound effects on PyC responses to non-occluded and occluded stimuli. In expert mice, the magnitude of responses to occluded stimuli was strongly increased (Fig. 2D-E), as was the percentage of responsive neurons (from 4% to 10%). In contrast, responses to non-occluded stimuli showed the opposite pattern. They weakened in expert mice, and fewer neurons were responsive (from 30% to 14%). During task performance, the percentage of neurons responsive to occluded stimuli was even higher (18%), while the decrease in the percentage of neurons responsive to non-occluded was less pronounced (21%).

As locomotion is known to modulate visual responses in mouse V1^18,30^, we assessed whether changes in locomotion influenced the observed plasticity of responses to non-occluded and occluded stimuli but found that this was not the case (Fig. S3A-B). We also ruled out the possibility that mice made eye movements in response to stimulus presentations, which might otherwise have positioned non-occluded image regions in the cRFs (Fig. S3C-D).

We conclude that visual experience decreased responses of L2/3 PyCs to non-occluded images, while it increased responses to the visual context provided by occluded images. Task engagement further enhanced responses to occluded images, suggesting that it strengthens responses to the visual contextual inputs. The observation that locomotion and task engagement did not significantly change the reduction of non-occluded image responses suggests that it is not caused by diminished attention.

### Separation of L2/3 PyCs responding to non-occluded and occluded images

We next analyzed how responses to non-occluded and occluded images were distributed across neurons. Interestingly, many PyCs appeared to respond to either the non-occluded stimuli or the occluded stimuli but not to both. This separation strengthened after familiarization with the images (“expert”) and was most pronounced in the condition under which the mice had been familiarized (“task”) (Fig. 2C, F, for responses to pairs of individual occluded and/or non-occluded images see Figs. S4 and S5). To quantify the extent of this unexpected separation of PyCs into two non-overlapping populations, we computed a separation index (SI) ranging from –1 to 1 (see Materials and Methods). The SI is positive when the degree of separation is higher than expected under the assumption that activity elicited by non-occluded and occluded stimuli is uncorrelated across PyCs. The SI is negative when both categories of stimuli activate the same neurons. The SI increased with experience and task engagement (Fig. 2F, SI: 0.040 for naive, 0.426 for expert and 0.874 for task). In parallel, the correlation coefficient (R) between population responses to non-occluded and occluded stimuli (averaged across images per neuron) decreased (R: 0.139 for naive, –0.036 for expert, –0.111 for task), affirming the separation in responses to the two types of stimuli.

The responses of PyCs to occluded images were reminiscent of inverse receptive field responses, which have been shown to depend on feedback inputs from higher order visual areas^31,32^. As responses to feedback inputs are characterized by a delay in response onset^27,31,32^, the onset latency of calcium responses to occluded images is expected to be longer than to non-occluded images if they are driven by feedback inputs. We determined latencies by fitting a curve to the activity time course and taking the time point at which the curve reached 33% of its maximum^32^.

We only included neurons with high quality fits (see Materials and Methods). Although calcium activity rises slower than spiking activity, the responses elicited by occluded stimuli were indeed delayed compared to responses elicited by non-occluded images (Fig. 2G, 189 ± 5.6 ms for non-occluded and 253 ± 8.0 ms for occluded, *P* < 0.001, LMEM). This delayed response is in line with the interpretation that they were predominantly driven by feedback inputs, but does not rule out a contribution of lateral connections within V1.

We tested whether the responses to occluded stimuli could be explained by responsiveness to the nearby edge of the occluded region, in which case the responsiveness would depend on proximity to this edge. However, the correlation between response strength and cRF distance from the edge of the occluded region was weak and statistically non-significant (Fig. S6, R = – 0.005, *P* = 0.876), consistent with findings from human fMRI and monkey recordings^23,27^.

After familiarization with non-occluded natural images, PyCs became more selective for individual images (Fig. 2H; lifetime sparseness). This increase in selectivity was associated with preferential suppression of neurons that responded to multiple images, as stimulus selectivity in naive mice was negatively correlated with the change in response amplitude after familiarization (R = −0.299, P = 0.001; Fig. S7). Selectivity for occluded images was much lower and did not increase with experience (Fig. 2H).

Taken together, our results indicate that responses to feedforward and contextual inputs predominantly occur in separate populations of L2/3 PyCs. Moreover, familiarization weakens responses to stimuli that included feedforward inputs, mostly affecting PyCs responding to multiple images. In contrast, responses to contextual inputs increase upon familiarization, in line with previous work^33^.

### Plasticity of responses to non-occluded and occluded images occurs in opposite directions in individual neurons

To understand how these changes at the population level were caused by changes in response properties of individual neurons, we matched neurons recorded at the different stages of the experiment and assessed how their responses to non-occluded and occluded images changed. This confirmed that PyCs responding to non-occluded images in naive mice mostly became less responsive in expert mice (Fig. S8) and when engaged in the task (Fig. 2I: yellow cells; Fig. 2J & S9: cells 1 & 5).

In contrast, a subset of PyCs that were initially unresponsive became responsive to occluded images after familiarization (Fig. 2I: red cells; Fig. 2J & S9: cells 2 & 3). This was also true for some PyCs that were initially responsive to non-occluded images (Fig. 2J & S9, cell 7). A small subset of neurons moved in the opposite direction of the majority of neurons (Fig. 2I & Fig. S8: blue and green cells; Fig. 2J & S9: cell 4). Interestingly, cells that increased their responsiveness to non-occluded images were highly selective for the individual familiarized images (lifetime sparseness of 0.48 in task, 0.69 in expert).

We conclude that also at the level of individual L2/3 PyCs, repeated visual stimulation drives plasticity of responses to contextual inputs alone in the opposite direction of those to stimuli that include feedforward inputs. This reorganization can shift cells from unresponsive or non-occluded image-driven toward strongly occluded image-driven. In a small subset of PyCs, however, changes occurred in the opposite directions.

### L2/3 PyCs responding to occluded images are not selective for large stimuli

We sought to determine why neurons responsive to occluded images do not respond strongly to non-occluded images, despite the presence of identical contextual stimuli. One explanation for this separation of responses could be that neurons responding to occluded images preferred large stimuli and responded to the occluder or its high-contrast edge, despite the absence of any change in the cRF input. However, the cRF sizes of PyCs responding to occluded or non-occluded images were similar (non-occluded: 17.16 ± 0.54°, occluded: 17.01 ± 0.67°, P=0.98, LMEM), which does not support this hypothesis. Moreover, as described above, responses were also not explained by proximity to the occluder edge (Fig. S6).

As a more direct test for selectivity for the occluder or its high contrast borders, we repeated the experiment with additional stimuli, including black and white noise stimuli – lacking the structure of natural images – with or without a gray or white occluder. To accelerate learning, we slightly changed the paradigm. The mice were now familiarized only by passive viewing of 4 of the 6 non-occluded images during 6 sessions across 3 days without rewards or lick training (Figure 3A). After familiarization, we again observed that distinct L2/3 PyC populations responded to occluded and non-occluded images (Figure 3B). Familiarization weakened responses to non-occluded images, while responses to occluded images were increased, albeit less strongly than after reward-assisted training (Figure 3C).

**Figure 3.**
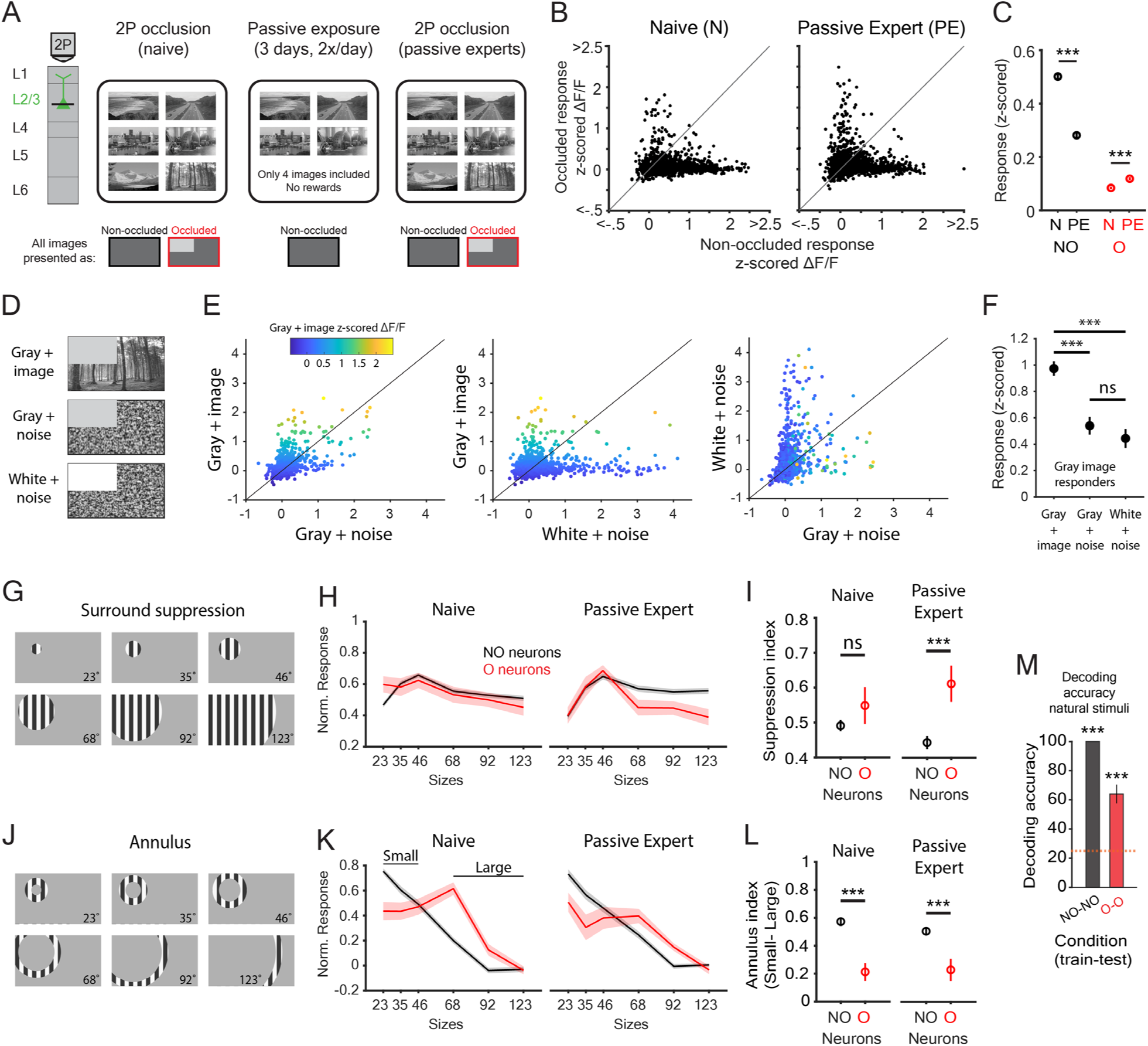
Response properties of L2/3 neurons responding to NO and O images. A) Schematic of experimental setup and passive exposure training regime. Mice passively viewed 4 of the 6 non-occluded images during 6 sessions in 3 days without lick training or rewards. B) Scatter plot showing average z-scored ΔF/F responses (t = 0.2-1 s) to the non-occluded (x-axis) and occluded (y-axis) stimulus across conditions. N = 1820 neurons (Naive) and 2106 neurons (Passive Expert) from 7 mice. C) Mean response strength across stimuli and conditions (mean ± SEM, averaged between t = 0.2-1 s). LMEM for all comparisons, ***: p<0.001. D) Example stimuli used for this experiment. Image + gray occl: familiar natural images with a gray occluder. Noise + gray occl: images consisting of black and white random noise with a gray occluder. Noise + white occl: images consisting of black and white random noise with a white occluder. E) Responses of all neurons in passive expert mice to the different stimuli in D), color coded by their response strength to familiar natural images with a gray occluder (image + gray occl response strength). F) Response strengths elicited by the different occluded stimuli, of neurons responsive to the “image + gray occl” condition (defined by response > 0.5 in E)). N = 99 neurons. LMEM for all comparisons, ***: p<0.001, ns: not significant. G) Surround suppression stimuli. The size in visual degrees has been indicated. H) Normalized responses of neurons responding to NO or O stimuli (defined by a response > 0.5). Note the stronger surround suppression for neurons responding to O stimuli. Tuning curves of each neuron were normalized to their maximum responses. Traces represent mean ± SEM over neurons. I) Mean surround suppression index for NO vs O responders in Naive and Passive Expert mice (mean ± SEM, calculated as (R_max_ – R_large_) / R_max_). NO neurons Naive: N = 691, O neurons Naive: N = 61. NO neurons Passive Expert: N = 392, O neurons Passive Expert: N = 100, from 7 mice. J) Annulus stimuli with their respective sizes in visual degrees as used in this experiment. The inner circle of the annulus matched the diameter of the surround suppression stimuli as shown in G). K) Normalized annulus response curves for neurons responding to NO or O stimuli (defined by response > 0.5). Note the stronger annulus responses at larger sizes for neurons responding to O stimuli. Tuning curves of each neuron were normalized to their maximum responses. Traces represent mean ± SEM over neurons. L) Mean annulus index for NO vs O responders in Naive and Passive Expert mice (mean ± SEM, calculated as R_small_ – R_large_). NO neurons Naive: N = 691, O neurons Naive: N = 61. NO neurons Passive Expert: N = 392, O neurons Passive Expert: N = 100, from 7 mice. M) Accuracy of population decoding of stimulus identity. An LDA decoder was trained on separate sets of trials for cross-validation of non-occluded responses (NO-NO, train-test) and occluded responses (O-O). Bars show mean ± SEM across repetitions of decoding runs. Chance performance (orange dashed line) was 25%. N = 2106 neurons from 7 mice. Permutation test, ***: p<0.001.

We then assessed whether L2/3 PyCs responsive to occluded natural images also responded to noise stimuli with gray or white occluders (Figure 3D). While some L2/3 PyCs responded to both occluded natural images and occluded noise stimuli, most L2/3 PyCs that responded to occluded natural images responded weakly to noise stimuli with a gray occluder and even less to those with a white occluder (Figure 3E). As a result, the average responses of L2/3 PyCs that were responsive to occluded natural images were markedly lower when stimulated with noise containing gray or white occluders (Figure 3F).

While these results do not rule out that a subset of PyCs responsive to occluded images are tuned to the grey occluder itself, they indicate that most of these neurons respond to ecRF inputs originating beyond the occluded region.

### Separation of L2/3 PyCs responding to occluded and non-occluded images is caused by differences in extraclassical receptive field inputs and surround suppression

We then tested another explanation for the separation of PyCs responding to non-occluded and occluded images. We reasoned that PyCs responding to occluded images may receive strong ecRF inputs, while being suppressed during presentation of non-occluded images. Such suppression could arise from local inhibitory mechanisms engaged when nearby PyCs with overlapping or adjacent cRF are co-activated, a phenomenon known as surround suppression (Fig. S10). Importantly, this form of suppression would be minimal for occluded images, as well as for the sparse noise stimuli used to map cRFs.

To test this hypothesis, we presented gratings in the form of discs and annuli of varying sizes to assess surround suppression and responses to stimuli beyond their cRFs, respectively, both before and after familiarizing mice with the non-occluded natural images.

We found that after familiarization, neurons responsive to occluded images showed significantly stronger surround suppression than those responsive to non-occluded images (Fig. 3G-I). Moreover, occluded-image responders showed weaker absolute responses to larger discs compared to non-occluded-image responders, both before familiarization (0.30 ± 0.044 vs. 0.41 ± 0.012) and even more prominently after familiarization (0.21 ± 0.025 vs. 0.45 ± 0.018). These results support the idea that neurons responsive to occluded images are more strongly suppressed by large stimuli.

Furthermore, neurons responsive to occluded images preferred larger annuli than those responsive to non-occluded images, and unlike the latter, they also responded to annuli positioned entirely outside the occluded quadrant (92°; Fig. 3J-L). This suggests that although occluded image-responders also received ecRF input from within the upper left quadrant, inputs from outside this region were sufficient to drive their responses to occluded images.

To assess whether responses to occluded images carried stimulus-specific information, we performed a population classification analysis using all neurons with cRFs within the occluded region, averaging responses for each neuron and each trial (between 0.2-1 s after stimulus onset). We used 50% of the trials to train a linear classifier (linear discriminant analysis; LDA) and the 50% remaining trials for testing. Training and testing the classifier on either non-occluded (Train-Test, i.e. NO-NO) or occluded (O-O) responses allowed us to examine whether these signals contain stimulus-specific information (Fig. 3M). We found that the identity of non-occluded images could be decoded with 100% accuracy. Decoding accuracy of occluded images was lower (64%), but well above chance level (25%), confirming a limited degree of specificity of L2/3 PyCs responses to occluded images.

Together, these results support the hypothesis that responses to occluded images arise from a combination of weakly selective contextual inputs from beyond the occluder in combination with release from surround suppression, possibly signaling a salient omission of a probable feedforward input.

### Responses to novel, non-familiarized images

We also asked whether the salient presence of an improbable feedforward input would also evoke enhanced responses. We therefore examined how L2/3 PyCs responded to the two images the mice had not been familiarized with (Fig. 4 & S11, novel stimuli, based on the first dataset obtained from our reward-assisted paradigm). Unlike the weakening of responses to familiar images, the activity elicited by these “novel” images had increased after training (Fig. 4A-D) as had the percentage of responsive neurons (21% in naive, 33% in expert mice). Notably, there was also an increase in the response strength to occluded novel images (Fig. 4A-D) and in the percentage of responsive neurons (3% in naïve, 8% in expert mice).

**Figure 4.**
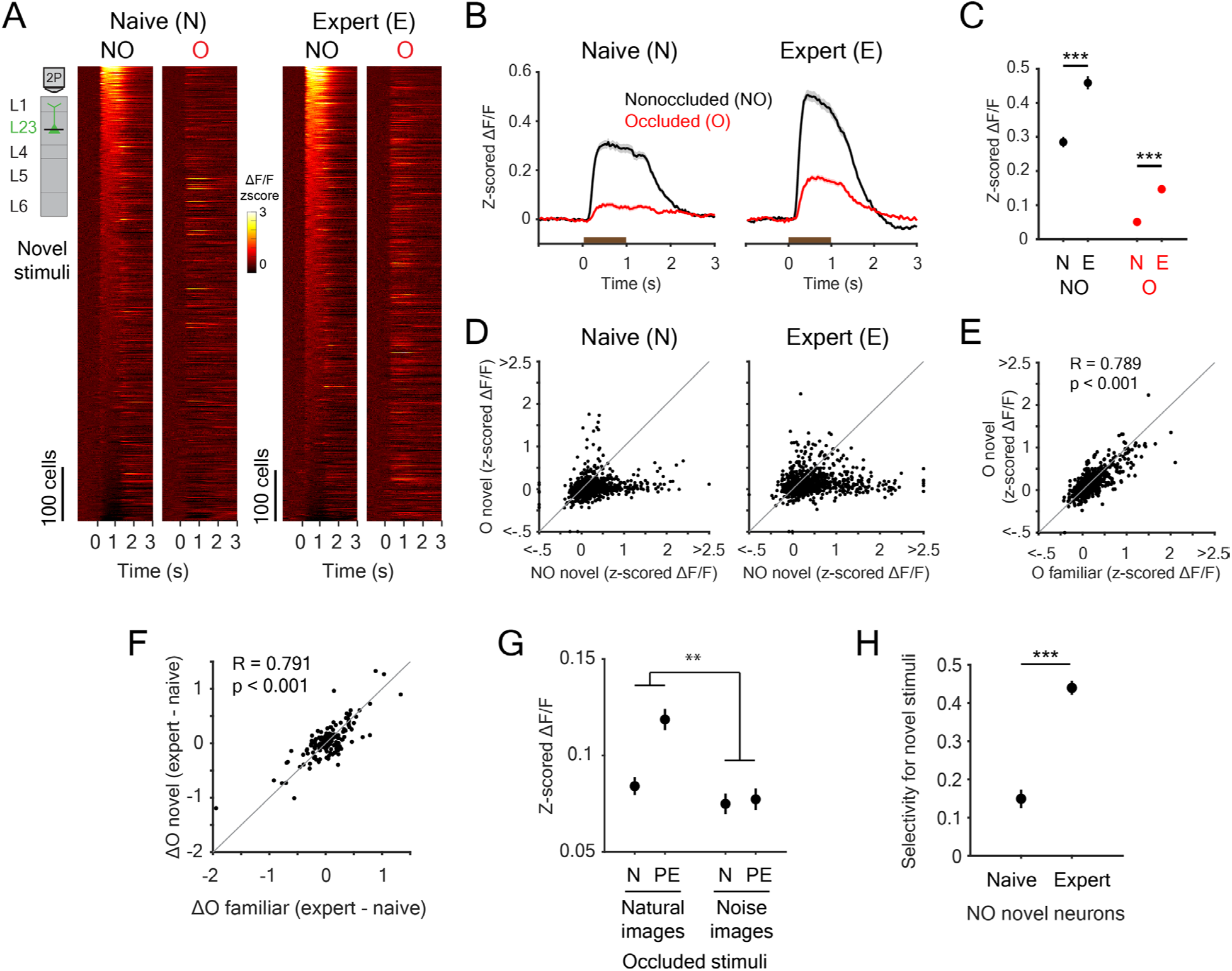
Novelty responses affect the most selective neurons and involve enhanced contextual inputs. A) Heatmap of z-scored ΔF/F responses for individual L2/3 neurons across conditions and stimuli for novel images. Neurons are sorted within each condition (N/E) based on their average response (t = 0.2-1 s) to the non-occluded stimuli. N = 841 neurons (Naive), 884 neurons (Expert) from 6 mice. B) Z-scored ΔF/F traces averaged across neurons and stimuli for novel images. Brown bars indicate the visual stimulus epoch. C) Mean response strength across stimuli and conditions (mean ± SEM, averaged between t = 0.2-1 s). LMEM for all comparisons, ***: p<0.001. D) Scatter plot showing average z-scored ΔF/F responses to non-occluded novel stimuli (x-axis) and occluded novel stimuli (y-axis) in naive mice (left) and expert mice (right). E) Scatter plot showing average z-scored ΔF/F responses to occluded familiar stimuli (x-axis) and occluded novel stimuli (y-axis) in expert mice. F) Scatter plot showing increase in average z-scored ΔF/F responses to occluded familiar stimuli (x-axis) and occluded novel stimuli (y-axis) of chronically matched neurons (N = 171 neurons). G) Mean response strength across training conditions to occluded stimuli as shown in Figure 3D. Familiarization increased responses to occluded versions of natural images, while it did not increase responses to occluded versions of noise images. **: p<0.01. N, Naïve; PE, Passive Expert. H) Selectivity for novel stimuli of responses to these stimuli in naive and expert mice. Included neurons responded to the novel NO image. Naive: N = 172 neurons, Expert: N = 285 neurons. Mean ± SEM, LMEM, p<0.001.

Given that responses to non-occluded images were more selective than responses to occluded images (Figs. 2H, 3J, S5B), we hypothesized that plasticity of these responses would follow the same pattern: repeated feedforward inputs would undergo selective weakening, whereas repeated ecRF inputs would be strengthened in a correspondingly broader, more generalized way.

This scenario yields three testable predictions. First, because novel images were not presented during training, responses selective for non-occluded novel images should not decrease. Second, if ecRF inputs strengthen in a way that generalizes across stimuli, the contextual inputs recruited by novel images should overlap with those recruited by familiar images. Such overlap could account for the enhanced responses to occluded novel images. Finally, because feedforward responses selective for non-occluded novel images remain intact, they could be further amplified by the strengthened contextual inputs, resulting in the increased responses observed for non-occluded novel images.

We found experimental support for all three predictions. First, we assessed the overlap in contextual inputs for familiar and novel images, by correlating the responses of PyCs to occluded versions of the familiar and novel images. PyCs that responded strongly to occluded novel images in expert mice indeed also responded to occluded familiar images (Fig. 4E, R=0.789, P<0.001). Moreover, neurons that increased their responses to the occluded novel images also increased their responses to occluded familiar images (Fig. 4F, R=0.791, P<0.001). One could argue that enhanced responses to occluded images reflect nonspecific surprise-driven amplification to any deviation from the familiarized stimulus. However, the occluded white noise stimuli in the second (passive familiarization) dataset did not show such enhancement after familiarization (Fig. 4G), despite the fact that they strongly deviate from the familiarized images. As the occluded white noise stimuli lack the structure of natural images, this finding instead supports the interpretation that ecRF responses strengthen and generalize across natural images sharing contextual information.

Second, we tested whether familiarization left responses to non-occluded novel images selectively intact. This is expected to increase selectivity for novel versus familiar images after familiarization. Indeed, we observed a significant experience-dependent increase in selectivity for novel over familiar images (Fig. 4H), consistent with a selective weakening of feedforward inputs.

Finally, if strengthened ecRF inputs amplify responses to novel images, PyCs responding to novel images should be more strongly influenced by these ecRF inputs. To test this, we quantified the relative contribution of visual input from inside versus outside the compound receptive field area (RFs of all neurons together) of the recorded neurons. We presented 3,600 non-occluded natural images to the mice that had been previously familiarized with four non-occluded images through passive exposure. For each neuron, we computed reverse correlations between local image structure and neural responses across this image set. As a measure of local image structure, we used Shannon entropy, which captures the spatial complexity of pixel intensities within a region. This approach allowed us to weight each image location by how strongly its local structure elicited neuronal responses.

Using these weights, we generated entropy-based activation maps for each neuron, indicating the spatial distribution of image features driving its responses. We then averaged these maps across neurons grouped by response type (occluded vs. non-occluded; familiar vs. novel). Each resulting map therefore reflects, for a given neuronal population, the spatial locations whose local image entropy most strongly contributed to their responses (Fig. S12A). By comparing the ratio of activity elicited by inputs outside versus within the compound cRF region across neuronal categories, we found that PyCs responding to novel non-occluded images received stronger ecRF inputs than those responding to familiar images (Fig. S12B).

Together, these findings support the idea that enhanced novelty responses in L2/3 PyCs arise from non-adapted selective responses to visual stimuli within the cRF, amplified by strengthened ecRF inputs that generalize across different natural scenes. This mechanism is consistent with the idea that interactions between feedforward and feedback inputs can generate mismatch-like responses^34^ but adds that opposing plasticity of these pathways is required to tune their interactions towards selective experience-based amplification of mismatched inputs.

### Non-occluded and occluded scenes elicited stimulus-specific neural signatures across species

Consistent with previous findings in humans and non-human primates, stimulus-specific activity in mouse V1 allowed reliable decoding of natural image identity from responses to both non-occluded and occluded stimuli^23,25,27^. We trained linear classifiers on population responses of PyCs (during task, using data from the first experiment) and tested decoding performance for non-occluded images (Train-Test, NO–NO), occluded images (O–O), and across conditions (NO– O and O–NO; Fig. 5A).

**Figure 5.**
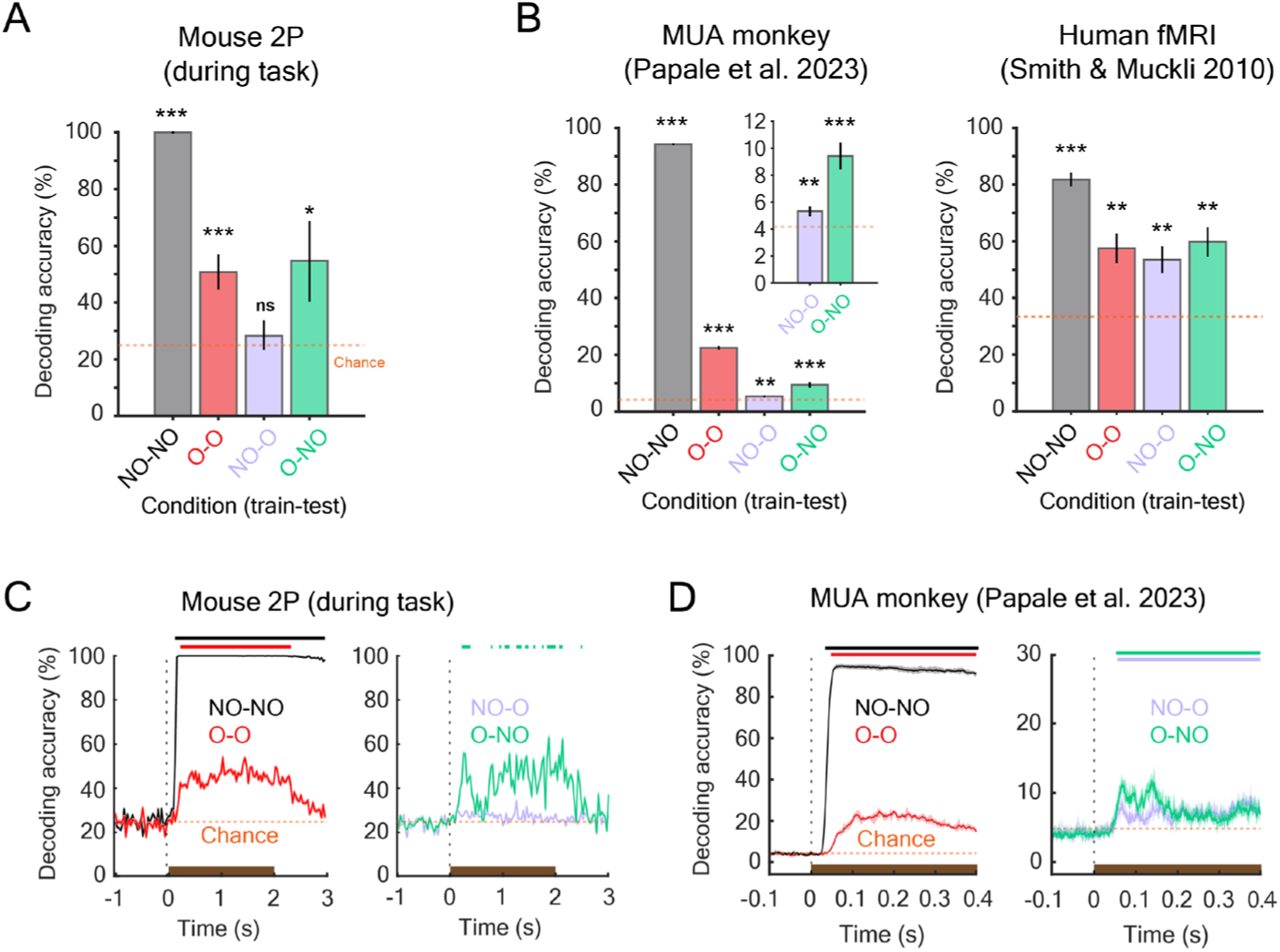
Responses to occluded scenes contain stimulus-specific information in mouse, monkey and human visual cortex. A) Accuracy of population decoding of stimulus identity for mouse PyCs (left, n = 300 PyCs) during the task. An LDA decoder was trained and tested on separate sets of trials for cross-validation (NO-NO, O-O) or cross-decoding (NO-O, O-NO). Bars show mean ± SEM across repetitions of decoding runs. Chance performance (orange dashed line) was 25%. Permutation test, ***: p<0.001, **: p<0.01, *: p<0.05, ns: not significant. NO: non-occluded, O: occluded. B) Left: decoding accuracy for multi-unit activity (MUA) monkey data. Chance performance was 4.16%. Permutation test, ***: p<0.001, **: p<0.01. Bars show mean ± SEM across repetitions of decoding runs. Inset shows a zoom for NO-O and O-NO conditions. Right: decoding accuracy for human fMRI data. Chance performance was 33.3%. Bars show mean ± SEM across individual subjects (n = 6). T-test against chance, ***: p<0.001, **: p<0.01. C) Population decoding accuracy over time for mouse PyCs. Colored dashes at the top depict significant time points (p<0.05, permutation test). Brown bars indicate the time window with the visual stimulus. Orange dashed line represents chance performance. D) As in C, but for monkey MUA data.

Decoding performance across conditions in mice was strikingly similar to that obtained by applying the same analysis to previously acquired data in humans and non-human primates (Fig. 5B-D), despite differences in species and recording techniques. Across all species and recording modalities, decoding accuracy was highest within stimulus conditions and lower for cross-condition decoding. We observed high (>80%) and significant decoding of responses to non-occluded stimuli (NO-NO). O-O decoding accuracy was lower but significant, just like in humans and non-human primates (Fig. 5B). Moreover, across all species and conditions, classifiers trained on non-occluded responses and tested on occluded responses (NO-O) were less accurate than the other way around. This asymmetry may reflect that classifiers trained on non-occluded images rely more strongly on feedforward responses, which are absent for occluded images, whereas classifiers trained on occluded images exploit contextual responses that also weakly contribute to non-occluded responses. Interestingly, the (O-NO) cross-decoding in mice only became significantly higher than chance during task (Fig. S13), suggesting that experience and task-engagement influenced the decoding efficiency.

The overall weaker cross-decoding in both directions, compared to O-O and NO-NO decoding, is not surprising given that responses to direct and contextual inputs arose predominantly from distinct neuronal populations. This limited cross-decoding may reflect a subset of neurons that responded to both full and occluded images – either weakly under both conditions, or strongly under one and weakly under the other. Such neurons may act as a bridge between direct and contextual representations, enabling partial generalization across these stimulus types.

### A computational model based on opposing plasticity rules in pyramidal cells reproduces dominant modes of experience-dependent plasticity

Based on these data, we hypothesized that two plasticity mechanisms with opposite directionality in individual PyCs enhance cortical responses to salient visual stimuli: selective weakening of responses to feedforward inputs and strengthening of contextual inputs that generalize across scenes. This strengthening amplifies responses of PyCs responsive to novel, non-adapted feedforward inputs and, if strong enough, enables responses by contextual inputs alone, when feedforward drive that recruits inhibition is absent (Fig. 6).

**Figure 6.**
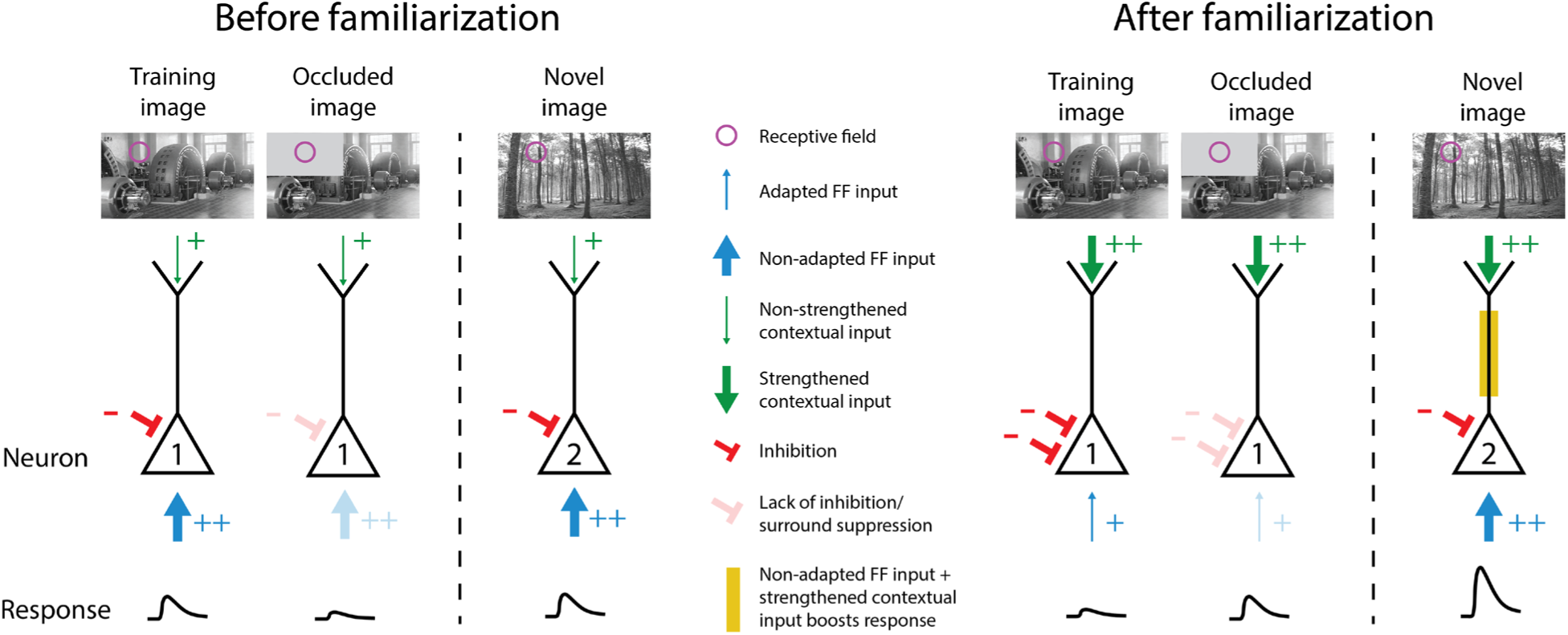
Opposing plasticity of feedforward and contextual inputs generates enhanced responses to salient stimuli. Before familiarization (left), PyCs integrate feedforward (FF) visual input within their classical receptive fields and contextual input from surrounding regions. Non-occluded training– and novel images (neurons 1&2) evoke responses driven through FF inputs and modulated by context. Occluded images elicit little activity due to weak contextual drive and absence of FF input (neuron 1). Upon familiarization (right), FF input from the training image selectively adapts and recruits stronger surround suppression (neuron 1), while contextual inputs strengthen in a generalizing way (neurons 1&2). As a result, responses to the non-occluded training image are reduced. In contrast, the occluded training image engages the strengthened contextual input in the absence of FF-driven inhibition, driving neural activity (neuron 1). The novel image also recruits this strengthened generalized context that enhances responses in PyCs that receive selective, non-adapted FF inputs from this image (neuron 2). Together, these opposing plasticity processes selectively amplify improbable inputs or omissions by maximizing the contrast between feedforward and contextual signals in individual neurons.

To explore this hypothesis, we developed a simple rate-based computational model (Fig. 7) of two-compartment PyCs, in which contextual input modulates responses at low intensities but drives the neuron when sufficiently strong^16,35^. In this model, feedforward inputs were selective and underwent depression as a function of postsynaptic PyC activity, while contextual inputs overlapped across presented stimuli and strengthened with repeated presentation. Feedforward-driven inhibitory inputs strengthened via Hebbian plasticity. All plasticity was restricted to synaptic inputs onto individual PyCs.

**Fig. 7.**
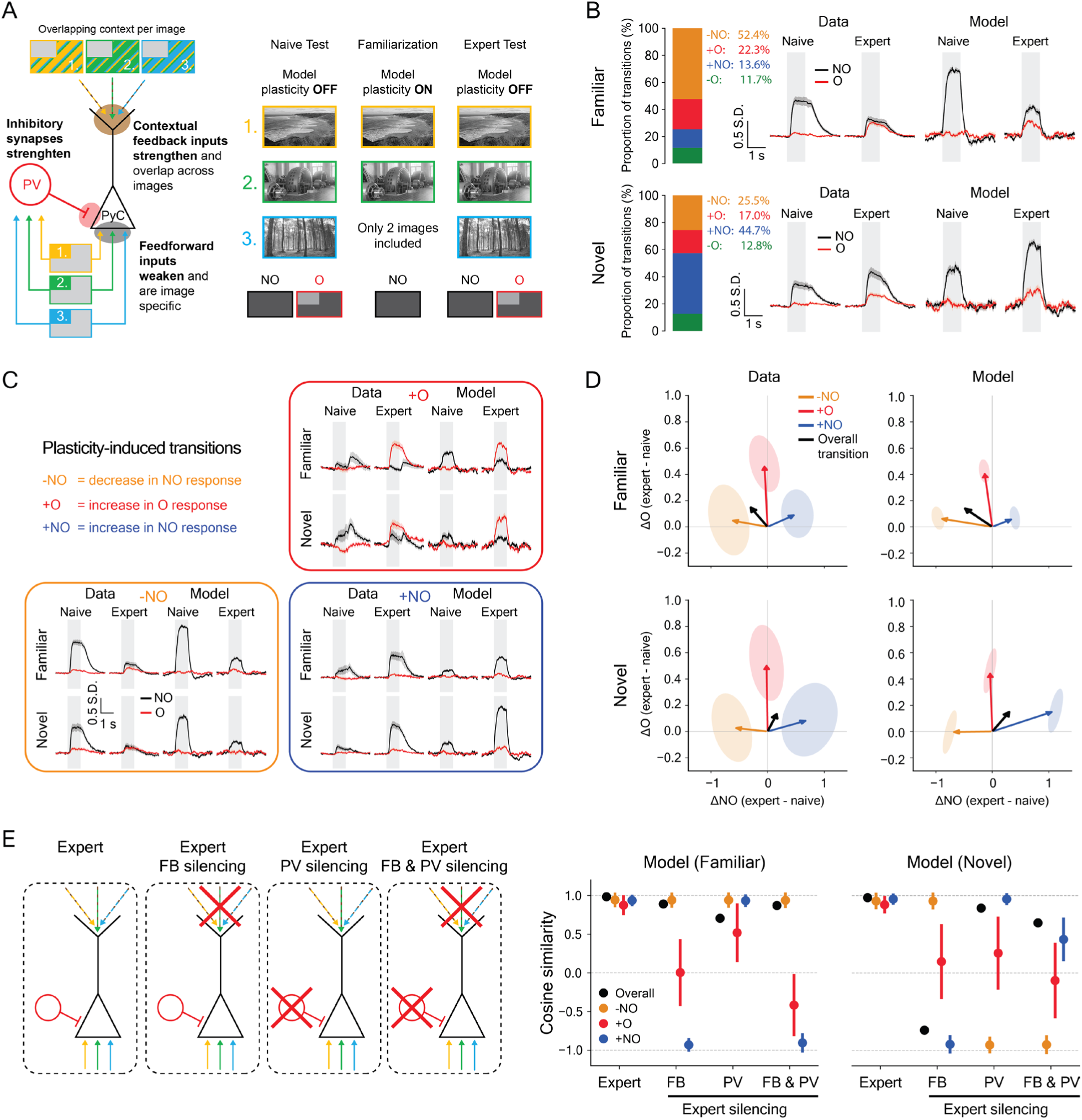
Computational model reproduces dominant modes of experience-dependent plasticity in L2/3 pyramidal cells. A) Model schematic and simulation setup. The PyC receives basal excitatory feedforward input, and apical excitatory contextual feedback (FB). The PV neuron receives excitatory feedforward input. Feedforward inhibition from PV acts as divisive normalization on PyC somatic activity which integrates basal and apical inputs. Only synapses onto PyC undergo plasticity. The simulation mirrors a simplified version of the real experimental setting by using two familiar inputs (1, 2) and one novel input (3). Each non-occluded (NO) image is represented as a combination of feedforward and contextual inputs, where feedforward inputs stimulate individual synapses while contextual inputs stimulate an overlapping set of apical synapses. We simulate occluded (O) images by zeroing-out the feedforward input for each image. Similar to the experiment, we perform naive testing on NO and O versions all inputs, but only familiarize with NO familiar inputs. B) Population-level response transitions to familiar and novel images. Response traces indicate overall population-level response transitions (± SEM), baseline-corrected and z-scored for both Data and Model. For data ± SEM over all recorded neurons. Proportion of individual recorded, chronically-matched neurons with ΔNO or ΔO > 0.3 undergoing different plasticity-induced response transitions to familiar and novel images. Population-level model response transition is obtained by a sum of responses of the –NO, +O, +NO models weighted by the real data proportions. C) Dominant modes of experience-dependent plasticity. Response traces indicate average over all recorded neurons undergoing either –NO, +O, or +NO transitions. For the model these response traces are averaged over 10 random seeds, for each of the six mean field models (separate parameter set fitted for each transition in familiar and novel). D) Transition vector representation of plasticity-induced transitions in the data and model. Within the ΔNO, ΔO plane, the direction and magnitude of a transition vector quantifies the direction and strength of plasticity-induced response changes to non-occluded and occluded images between naive and expert conditions. For Data, ellipsoids indicate covariance over transition vectors of individual recorded neurons. For Model, ellipsoids indicate covariance over different random seeds. The overall transition vector is a weighted sum of the three –NO, +O, +NO transition vectors, weighted by the proportion of those transitions in the real data, as reported in B). E) Model ablation experiments. After familiarization we performed expert testing on NO and O versions of all inputs; coupled with ablating PV and feedback. Cosine similarity (cosine of the angle between two vectors) plots show average cosine similarity (error bars show ± SD) of each of the six model’s transition vectors under a given ablation condition against naive-expert transition vectors of all recorded and chronically-matched neurons undergoing corresponding plasticity-induced transition.

We simulated familiarization with two non-occluded images and measured responses to familiar and novel images, with and without occlusion (Fig. 7A). Non-occluded images provided feedforward inputs to inhibitory cells and PyCs and contextual inputs to PyCs, while occluded images only provided contextual inputs to PyCs.

We modeled the three dominant transition types separately using the same model, varying a small number of parameters controlling feedforward and feedback input strengths, PV-mediated inhibition, and apical gain (Table S2): reduced responses to non-occluded images (−NO), enhanced responses to occluded images (+O), and enhanced responses to non-occluded images (+NO). To match the data, we modeled the −NO transition as stronger for familiar images and the +NO transition as stronger for novel images, while +O transition models were similar across conditions. Weighting the modeled transitions by their observed proportions in the data, with more −NO transitions for familiar and more +NO transitions for novel images, closely reproduced the population-average responses (Fig. 7B).

The same model thus reproduced reduced responses to familiar non-occluded images and enhanced responses to familiar occluded images, as well as enhanced responses to occluded and non-occluded novel images, depending on the initial conditions (Fig. 7C, D). To also account for the small population of highly selective PyCs whose responses to familiar non-occluded images increased (Fig. 2I, S8), we gated the rate of basal plasticity by a current slowly accumulating with repeated basal activity, resulting in weaker feedforward depression in cells responding to only one image (in line with Fig. S7). This allowed strengthened contextual inputs to outweigh mild feedforward adaptation, reproducing the +NO transition for familiar images (Fig. 7C, D).

The −NO, +O and +NO transitions were robust to variation in individual model parameters. For most parameters, transition direction and magnitude were largely preserved across the full tested range, from 0.05 to 5 times the fitted value, although some parameters showed narrower ranges of robustness (Fig. S14A-C). While the model thus reliably captured the most frequent transitions, it did not address the less common reduction in responses to occluded images (−O). Accounting for this transition may require extending the model to include lateral inhibition or plasticity in cortical areas providing contextual feedback.

Ablating contextual inputs after familiarization reduced responses to familiar and novel occluded images (Fig. 7E, S15), and reversed the +NO transitions in both conditions, indicating that these response increases depended on strengthened contextual inputs. Conversely, removing inhibition caused +O PyCs to respond strongly to non-occluded images, supporting a role for feedforward-driven inhibition in separating this response population from PyC responding to non-occluded images after familiarization (Fig. 7E, S15). Interestingly, −NO transitions persisted after either ablation for familiar images, consistent with depression of feedforward inputs, whereas −NO transitions for novel images reversed upon removing inhibition, suggesting a stronger contribution of inhibitory plasticity.

Together, these simulations support the idea that opposing plasticity of inputs onto individual PyCs, combined with feedforward-driven inhibition, provide a possible explanation for the principal familiarization-induced response changes.

## DISCUSSION

Our results show that familiarization to natural images induces a striking rebalancing of L2/3 PyCs responses in V1 to non-occluded and occluded images. Repeated responses to non-occluded images providing direct sensory inputs are selectively weakened, whereas contextual responses to occluded images strengthen and generalize across images. Acting together within individual neurons, these opposing plasticity mechanisms may enable PyCs to extract salient signals and omissions from predictable visual backgrounds, without requiring a difference computation.

Several mechanisms may contribute to the selective downscaling of neural responses to direct visual inputs after familiarization. These include synaptic depression^36,37^ and reduced intrinsic excitability that both occur in relatively short time spans^38,39^, and synaptic scaling via decreased postsynaptic AMPA receptor conductance over longer periods^40,41^. Additionally, repeated activation can strengthen inhibition, suppressing highly active excitatory neurons^42,43^ and contributing to surround suppression. Acting across different timescales and levels of organization, these mechanisms may synergistically reduce responses to familiar stimuli, while contextual influence is increasing.

Repeated presentation of natural visual scenes particularly reduced responses of less selective PyCs that responded to multiple images during familiarization. This disproportionate reduction of broadly-tuned neuron responses increased the relative contribution of more selective neurons, leading to a sharpening of visual responses, in accordance with previous work^44–46^. The observed sharpening is consistent with an adaptive normalization mechanism, in which gain reductions scale with a neuron’s cumulative activity during repeated stimulus exposure^47^. This sharpening is not expected in classical formulations of hierarchical predictive coding that emphasize feedback-driven inhibition ‘explaining away’ expected inputs, resulting in dampening^44,48–52^.

A subset of PyCs responded to occluded natural scenes despite the absence of cRF input, reminiscent of inverse receptive field responses driven by feedback inputs in the absence of feedforward drive^31^. This population of occluded image responders increased following familiarization with non-occluded scenes. Most of these neurons did not respond to the non-occluded familiar images, suggesting that strengthening of ecRF inputs does not require somatic spiking activity. This observation aligns with evidence that potentiation of feedback inputs onto L2/3 PyC dendrites can occur independently of somatic firing^53^ but may require dendritic NMDA spikes^54^. Disinhibition may have also supported the strengthening of ecRF inputs^32,55^. However, since most PyCs that responded to occluded images in expert mice did not at all respond to them before familiarization, disinhibition alone is unlikely to explain the observed changes. Possibly, disinhibition may have facilitated plasticity of ecRF inputs onto L2/3 pyramidal cells^56,57^.

While PyC responses to different occluded images generalized across scenes, they nevertheless carried image-specific information, as stimulus identity could be decoded from these responses. This observation is consistent with a recent report showing that responses to occluded regions generalize across scenes while still retaining stimulus-specific information^58^. Moreover, because PyCs responding to occluded images also received input from more proximal visual locations, as revealed by annulus stimuli, they appear to integrate spatially distributed contextual signals across the surround. Synaptic strengthening in apical dendrites depends on the co-activation of nearby synapses⁵⁰, suggesting that this form of plasticity could support the learning of structured contextual regularities rather than merely reinforcing frequently occurring features. Such contextual learning at the level of individual neurons may therefore enable selective amplification of inputs that violate these learned regularities. Future experiments will be needed to characterize the statistical structure of these contextual signals and to determine how it depends on stimulus proximity and spatial arrangement.

Occlusion-responsive neurons exhibited more surround suppression and received more distal ecRF inputs than those responding to non-occluded images. This combination of response properties may explain responses to occluded images, but also inverse receptive field responses^31,32^. In this scenario, weakly selective inhibition from local interneurons^59^, driven when neighboring neurons are activated by the non-occluded stimulus, suppresses occluded image responders. When feedforward input near the cRF is absent, inhibition is reduced, allowing input from the surrounding visual context to drive the PyC’s response (Fig. S10). This mechanism, requiring feedforward-driven inhibition, would also explain why responses to occluded and non-occluded images occur in separate neuronal population. These responses to occluded regions may highlight regions where visible input is missing from a scene, contributing to the perception of such areas as figures^32,58^. Whether such responses can also support contextual completion^60^ when parts of familiar scenes are blurred or ambiguous is currently unclear.

Strengthening of ecRF inputs may also explain enhanced responses elicited by novel images presented after familiarization. Because these contextual inputs generalized across familiarized and novel images, PyCs selective for these novel images likely receive stronger visual contextual inputs after familiarization. However, unlike PyCs responsive to familiar images, they did not undergo adaptation during familiarization. Consequently, the coincidence of non-adapted feedforward and strengthened contextual input may enhance feedforward drive in a context-dependent way^61^. This could involve apical amplification, consistent with models in which apical dendrites integrate contextual input and modulate somatic gain^62–65^. Supporting this idea, we observed that PyCs responding to novel images were more strongly influenced by inputs outside their cRFs than those responding to familiar images. It is well possible that additional novelty related inputs, such a thalamic or neuromodulatory inputs, further enhance responses to occluded or novel images^4^.

While most PyCs reduced their responses to familiar images, a small subset showed increased responsiveness and strong selectivity for familiarized stimuli. One explanation for this observation is that selective neurons may adapt less as they respond less frequently, and actually become more responsive due to the enhanced contextual inputs. We can indeed simulate this response strengthening of selective neurons by making them adapt more slowly. Alternatively, these neurons may not compute how feedforward inputs relate to context, but instead encode learned, behaviorally relevant signals. If so, they could represent a distinct subtype of L2/3 PyCs shaped by Hebbian plasticity, potentially enabled by a different source of feedback aligned with their cRF inputs. Consistent with this idea, it was found that L2/3 feedback aligns with cRF responses, whereas feedback from L5 PyCs in the lateromedial area originates from more distal ecRF regions and is not anatomically aligned with postsynaptic orientation preference^68^.

Recent work showed that the emergence of such mismatched feedback during development can be modeled by Hebbian plasticity of feedforward inputs combined with anti-Hebbian plasticity of feedback inputs^67^. While this developmental plasticity also wires PyCs to enhance responses to deviations between feedforward and feedback signals, its functional outcome differs from what we observe in adult V1: the model predicts that experienced feedforward responses are strengthened, whereas matching feedback connections become disconnected.

Notably, decoding of non-occluded and occluded images, as well as cross-decoding between them, was similarly efficient across humans, monkeys, and mice, despite being measured with fMRI, electrophysiology, and 2P calcium imaging, respectively. This convergence across species and recording modalities suggests that common neural mechanisms underlie contextual processing, consistent with the circuit principles described above. The above-chance decoding of occluded natural images in naive mice suggests that contextual inputs are already structured before familiarization with these images, potentially reflecting connectivity shaped during development or by natural vision during adulthood. Ongoing plasticity would then allow this connectivity to be continuously refined, tuning the system to the statistical regularities of the animal’s current environment and ensuring that saliency is assigned relative to what is presently familiar.

A simple circuit-level explanation consistent with our findings, is one in which two plasticity mechanisms with opposite directionality in individual PyCs dynamically optimize cortical responses to salient visual stimuli (Fig. 6). One mechanism selectively weakens responses to frequently active feedforward inputs^36,40,41,44,48,68–71^, while the other strengthens regularly active contextual inputs^33,53,54,56^ that generalize across scenes. This strengthening has two major consequences: 1) it amplifies responses to novel, non-adapted feedforward inputs that violate the statistical structure of familiarized contexts and 2) when local feedforward drive recruiting inhibition is absent, it enables contextual signals to drive responses that may signal the absence of expected inputs. By maximizing the contrast between feedforward and contextual influences within individual L2/3 PyCs, this circuitry provides a plausible cellular mechanism for detecting and enhancing salient information.

When we explored this circuit mechanism in a computational model, we found that simple plasticity rules operating exclusively at the cellular level were sufficient to reproduce the dominant response transitions, including the counterintuitive strengthening of image-selective responses to familiarized images. Feedback-driven inhibition was not required. Importantly, the model demonstrates how a single mechanism – strengthened contextual inputs – can account for enhanced responses to both novel and occluded images, through dendritic amplification and dendritic drive, respectively.

This context-contrasting mechanism fits well with Helmholtz’s classic idea of unconscious inference, in which perception reflects experience-based interpretation rather than passive sensory reception^72^. By amplifying responses to deviations from learned contextual structures, this circuitry may support rapid detection of behaviorally relevant stimuli within familiar scenes. When feedforward input is missing, strengthened context-driven responses may highlight such occluded regions, contributing to their perception as coherent shapes. These responses may also support contextual completion when inputs are degraded or ambiguous. Importantly, this concept may also help shed light on psychiatric symptoms such as psychosis: if the processes underlying context-contrasting plasticity are disrupted, the brain may assign saliency to inappropriate inputs, potentially contributing to hallucinations, delusions, or disorganized thinking^73^. By identifying a potential cellular mechanism for disrupted saliency processing within individual neurons, context-contrasting circuitry may offer new mechanistic approaches for understanding these conditions.

### Resource availability

Processed data per figure panel and MATLAB scripts for reproducing the figures will be available here: osf.io/n8w5m

Unprocessed two-photon imaging data are available from the lead contact upon request.

The code for the computational model is available here: https://github.com/Leveltlab/context-contrasting-model

## Acknowledgements

We thank all members of the Levelt lab for discussion and support. We thank staff of the animal facility and mechatronics department at the Netherlands Institute for Neuroscience for technical support. We thank Dr. Chris van der Togt, Dr. Enny van Beest, Paul Neering, Maaike van der Aa, Marija Žuvela, Jessie Lof, Augustijn Vrolijk and Anda de Witte for help with experiments and analyses. We thank Dr. William Phillips for the critical reading of the manuscript. We thank Dr. Fraser Smith for help with the human data. This project received funding from the European Union’s Horizon 2020 Research and Innovation Program under grant agreement nos. 785907 (HBP SGA2, CL, PR, ML & LM) and 945539 (HBP SGA3, CL, PR, ML & LM), from the Biotechnology and Biological Sciences Research Council (BBSRC) “Layer-specific cortical feedback (LM with LSP), NWO VENI n. VI.Veni.222.217 (PP), ERC Starting Grant n. 101219949 “SteerMEM” (PP), from the Friends Foundation of the Netherlands Institute for Neuroscience, made possible by a private endowment (OD, CNL), and from NWO OCENW.KLEIN.178 (CNL). The authors declare that they have no competing interests. All data needed to evaluate the conclusions in the paper are present in the paper and/or the Supplementary Materials.

## Author contributions

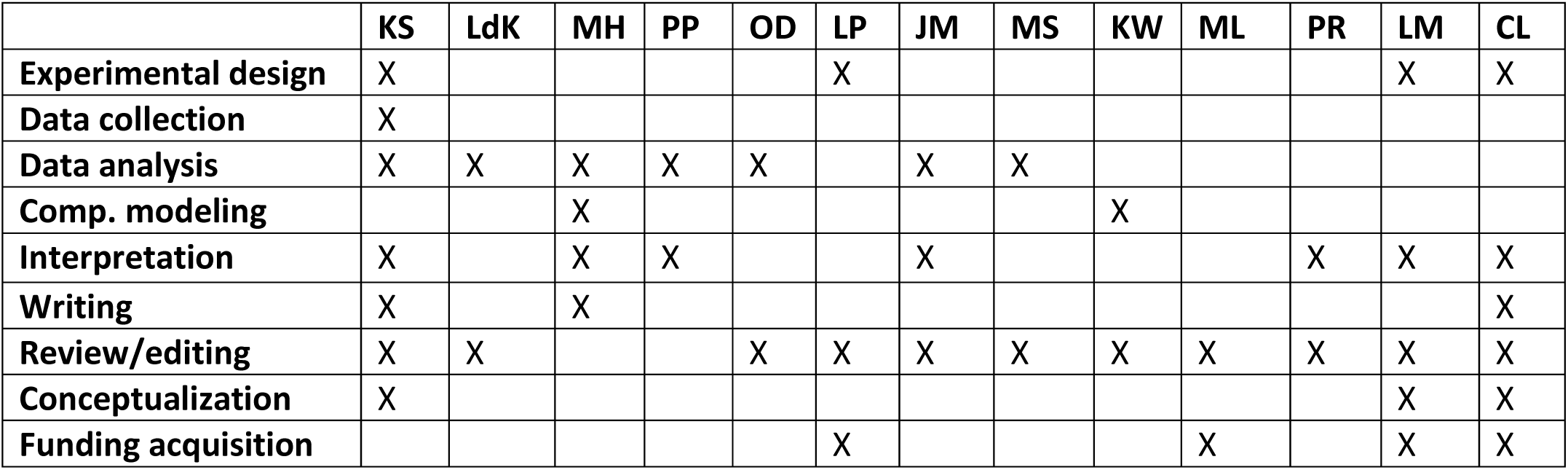

## Declaration of generative AI and AI-assisted technologies in the writing process

During the preparation of this manuscript, the authors used ChatGPT to assist with improving the readability of the text. The authors reviewed and edited all AI-assisted output and assume full responsibility for the final manuscript.

## Supplemental information

Supplemental Figures S1-S15

Supplemental Tables S1-S2

Statistical table.

Statistical table
| Figure | Comparison | Parameter | Test | # mice | N represents | # neurons | p-val | Sign |
| --- | --- | --- | --- | --- | --- | --- | --- | --- |
| **2E** | Naive vs Expert | NO response | LMEM-Anova | 6 vs 6 | Neurons | 841 vs 884 | <0.001 | *** |
|  | Naive vs Task | NO response | LMEM-Anova | 6 vs 6 | Neurons | 841 vs 300 | <0.001 | *** |
|  | Expert vs Task | NO response | LMEM-Anova | 6 vs 6 | Neurons | 884 vs 300 | 0.9999 | ns |
|  | Naive vs Expert | O response | LMEM-Anova | 6 vs 6 | Neurons | 841 vs 884 | <0.001 | *** |
|  | Naive vs Task | O response | LMEM-Anova | 6 vs 6 | Neurons | 841 vs 300 | <0.001 | *** |
|  | Expert vs Task | O response | LMEM-Anova | 6 vs 6 | Neurons | 884 vs 300 | <0.001 | *** |
| **2G** | NO vs O | Response onset | LMEM-Anova | 6 vs 6 | Neurons | 584 vs 528 | <0.001 | *** |
| **2H** | Naive vs Expert | NO selectivity | LMEM-Anova | 6 vs 6 | Neurons | 328 vs 223 | <0.001 | *** |
|  | Naive vs Task | NO selectivity | LMEM-Anova | 6 vs 6 | Neurons | 328 vs 79 | <0.001 | *** |
|  | Expert vs Task | NO selectivity | LMEM-Anova | 6 vs 6 | Neurons | 223 vs 79 | <0.001 | *** |
|  | Naive vs Expert | O selectivity | LMEM-Anova | 6 vs 6 | Neurons | 79 vs 182 | 0.9999 | ns |
|  | Naive vs Task | O selectivity | LMEM-Anova | 6 vs 6 | Neurons | 79 vs 79 | 0.9999 | ns |
|  | Expert vs Task | O selectivity | LMEM-Anova | 6 vs 6 | Neurons | 182 vs 79 | 0.9999 | ns |
| **3C** | N vs PE | NO response | LMEM-Anova | 7 vs 7 | Neurons | 1820 vs 2106 | <0.001 | *** |
|  | N vs PE | O response | LMEM-Anova | 7 vs 7 | Neurons | 1820 vs 2106 | <0.001 | *** |
| **3F** | NO cells vs O cells (N) | Surround suppression | LMEM-Anova | 7 vs 7 | Neurons | 691 vs 61 | 0.302 | ns |
|  | NO cells vs O cells (PE) | Surround suppression | LMEM-Anova | 7 vs 7 | Neurons | 392 vs 100 | <0.001 | *** |
| **3I** | NO cells vs O cells (N) | Annulus index | LMEM-Anova | 7 vs 7 | Neurons | 691 vs 61 | <0.001 | *** |
|  | NO cells vs O cells (PE) | Annulus index | LMEM-Anova | 7 vs 7 | Neurons | 392 vs 100 | <0.001 | *** |
| **3J** | NO-NO vs chance | Decoding accuracy | LMEM-Anova | 7 | Neurons | 2106 | <0.001 | *** |
|  | O-O vs chance | Decoding accuracy | LMEM-Anova | 7 | Neurons | 2106 | <0.001 | *** |
| **4C** | Naive vs Expert | NO response | LMEM-Anova | 6 vs 6 | Neurons | 841 vs 884 | <0.001 | *** |
|  | Naive vs Expert | O response | LMEM-Anova | 6 vs 6 | Neurons | 841 vs 884 | <0.001 | *** |
| **4G** | Naive vs Expert | NO selectivity | LMEM-Anova | 6 vs 6 | Neurons | 172 vs 285 | <0.001 | *** |
| **5A** | NO-NO vs chance | Decoding accuracy | Permutation test | 6 | Neurons | 300 | <0.001 | *** |
|  | O-O vs chance | Decoding accuracy | Permutation test | 6 | Neurons | 300 | 0.001 | ** |
|  | NO-O vs chance | Decoding accuracy | Permutation test | 6 | Neurons | 300 | 0.018 | * |
|  | O-NO vs chance | Decoding accuracy | Permutation test | 6 | Neurons | 300 | 0.001 | ** |
| **5B** | NO-NO vs chance | Decoding accuracy | Permutation test | 175 | Recording sites | NA | <0.001 | *** |
|  | O-O vs chance | Decoding accuracy | Permutation test | 175 | Recording sites | NA | <0.001 | *** |
|  | NO-O vs chance | Decoding accuracy | Permutation test | 175 | Recording sites | NA | 0.007 | ** |
|  | O-NO vs chance | Decoding accuracy | Permutation test | 175 | Recording sites | NA | <0.001 | *** |
|  | NO-NO vs chance | Decoding accuracy | t-test | 6 | Humans | NA | <0.001 | *** |
|  | O-O vs chance | Decoding accuracy | t-test | 6 | Humans | NA | 0.0058 | ** |
|  | NO-O vs chance | Decoding accuracy | t-test | 6 | Humans | NA | 0.0081 | ** |
|  | O-NO vs chance | Decoding accuracy | t-test | 6 | Humans | NA | 0.0038 | ** |
| **S2A** | Naive vs Expert | NO response | LMEM-Anova | 6 vs 6 | Neurons | 884 vs 841 | <0.001 | *** |
|  | Naive vs Task | NO response | LMEM-Anova | 6 vs 6 | Neurons | 884 vs 300 | <0.001 | *** |
|  | Expert vs Task | NO response | LMEM-Anova | 6 vs 6 | Neurons | 841 vs 300 | 0.9999 | ns |
|  | Naive vs Expert | O response | LMEM-Anova | 6 vs 6 | Neurons | 884 vs 841 | <0.001 | *** |
|  | Naive vs Task | O response | LMEM-Anova | 6 vs 6 | Neurons | 884 vs 300 | <0.001 | *** |
|  | Expert vs Task | O response | LMEM-Anova | 6 vs 6 | Neurons | 841 vs 300 | <0.001 | *** |
| **S2B** | Naive vs Expert | NO response | LMEM-Anova | 6 vs 6 | Neurons | 884 vs 841 | <0.001 | *** |
|  | Naive vs Task | NO response | LMEM-Anova | 6 vs 6 | Neurons | 884 vs 300 | 0.0333 | * |
|  | Expert vs Task | NO response | LMEM-Anova | 6 vs 6 | Neurons | 841 vs 300 | 0.2348 | ns |
|  | Naive vs Expert | O response | LMEM-Anova | 6 vs 6 | Neurons | 884 vs 841 | <0.001 | *** |
|  | Naive vs Task | O response | LMEM-Anova | 6 vs 6 | Neurons | 884 vs 300 | <0.001 | *** |
|  | Expert vs Task | O response | LMEM-Anova | 6 vs 6 | Neurons | 841 vs 300 | <0.001 | *** |
| **S2D** | Position vs zero, NO, X | Pupil position | Signrank | 6 vs 6 | Mice | NA | 0.6875 | ns |
|  | Position vs zero, NO, Y | Pupil position | Signrank | 6 vs 6 | Mice | NA | 0.9999 | ns |
|  | Position vs zero, O, X | Pupil position | Signrank | 6 vs 6 | Mice | NA | 0.5625 | ns |
|  | Position vs zero, O, Y | Pupil position | Signrank | 6 vs 6 | Mice | NA | 0.3125 | ns |
| **S9B** | NO fam vs O fam | Activity ratio | Permutation test | 6 vs 6 | Neurons | 277 vs 63 | 0.0060 | ** |
|  | NO fam vs NO nov | Activity ratio | Permutation test | 6 vs 6 | Neurons | 277 vs 433 | 0.0060 | ** |
|  | NO fam vs O nov | Activity ratio | Permutation test | 6 vs 6 | Neurons | 277 vs 52 | 0.0060 | ** |
|  | O fam vs NO nov | Activity ratio | Permutation test | 6 vs 6 | Neurons | 63 vs 433 | 0.0060 | ** |
|  | NO nov vs O nov | Activity ratio | Permutation test | 6 vs 6 | Neurons | 433 vs 52 | 0.0060 | ** |
|  | O fam vs O nov | Activity ratio | Permutation test | 6 vs 6 | Neurons | 63 vs 52 | 0.1080 | ns |

## MATERIALS AND METHODS

### Mice

All experiments were approved by the institutional animal care and use committee of the Royal Netherlands Academy of Arts and Sciences. We used 4 female and 2 male G35-3Dtl^28^ crossed to AI93 TITL-GCaMP6f mice (Jackson Laboratories, www.jaxmice.jax.org, strain 024107). Mice were group housed under a 12 h reversed day/night cycle and provided with ad libitum access to food and water. Experiments were performed in the dark phase. Experimental animals were adult at surgery (>40 days) and both males and females were used. Mice had access to a running wheel in their home cage throughout the experiment.

### Window surgery

For the cranial window surgery, mice were anesthetized with isoflurane (4% induction, 1.6% maintenance in oxygen). During the surgeries, temperature was maintained at 37°C with euthermic pads and eyes were protected from light and drying using Cavasan eye ointment. Mice were implanted with a double (3+4 mm diameter) glass window and a custom metal head-ring on the center of V1 to allow head-fixation. The glass window and head-ring were fixed to the skull using dental cement. At least one week after the window surgery, mice were handled and trained to be head immobilized in our setup until they sat comfortably and started running regularly on their own.

### Two-photon imaging

Two-photon experiments were performed on a two-photon microscope (Neurolabware) equipped with a Ti-sapphire laser (Mai-Tai ‘Deepsee’, Spectraphysics; wavelength, 920 nm) and a 16x, 0.8 NA water immersion objective (Nikon) at a zoom of 1.6x. The microscope was controlled by Scanbox (Neurolabware) running on MATLAB (Mathworks). Images were acquired at a frame rate of 15.5 (RF mapping) or 31 Hz (occlusion experiments) in V1 L2/3 at a depth of 150-250 µm from the cortical surface. For all behavioral training and imaging sessions, mice were head restrained and free to run on a 3D-printed running wheel (8.5 cm diameter). Locomotion speed was measured with a rotary encoder and processed using Arduino and MATLAB.

All visual stimuli for two-photon imaging were presented on a 24-inch gamma-corrected non-occluded HD LED monitor (1920×1080 pixels) using OpenGL and Psychophysics Toolbox 3 running on MATLAB. The monitor was positioned centered at 15 cm away from the mouse, yielding a field-of-view of 120° x 90°.

For RF mapping, we used eccentricity-corrected sparse noise stimuli with squares of 7.5 visual degrees at different locations covering the entire screen. Each trial consisted of 3 black and 3 white randomly positioned squares on a gray background. Stimuli were presented for 0.5 s with a gray screen ITI of 1.5 s.

For occlusion experiments, we presented 6 natural images that were part of a dataset used for similar experiments before^23,27^. The images were presented in gray scale and were corrected for overall luminance across images. We presented images non-occluded or partially occluded using a gray occluder positioned on the top left quadrant of the screen. The occluder had identical luminance to the ITI gray screen. Non-occluded and occluded images (50% of trials each) were both presented 20 times for 1 s with a random ITI of 4-6 s.

For surround suppression experiments, we presented circular static black and white gratings (0.05 cycles per degree, 90% contrast) of 4 orientations (0°, 45°, 90° and 135°) and 6 sizes (diameter of 23°, 35°, 46°, 68°, 92° and 123°) at the location of the screen that represented the aggregate RF location of the entire imaging field-of-view. Stimuli were presented 10x each for 1 s with a random ITI of 4-6 s.

For annulus experiments, we presented circular static black and white rings (spatial frequency: 0.05 cycles per degree, 90% contrast) of 4 orientations (0°, 45°, 90° and 135°) and 6 sizes (inner diameter of 23°, 35°, 46°, 68°, 92° and 123°) at the location of the screen that represented the aggregate RF location of the entire imaging field-of-view. The ring width was 4 cm. Stimuli were presented 10x each for 1 s with a random ITI of 4-6 s.

For noise stimuli with gray or white occluders we presented images of random black and white smoothed noise with a mean luminance that matched the ITI gray screen. We presented these images with gray and white occluders. Stimuli were presented 10x each for 1 s with a random ITI of 4-6 s.

To estimate the influence of cRF and ecRF information on familiar and novel image responding PyCS, we presented 4000 images from 720 classes taken from the THINGS database^74^. Images were shown for 0.5 s, followed by a delay of 0.5 s where mice viewed a gray screen.

### Wide-field imaging

Wide-field imaging was performed as described as before^75^. Briefly, we imaged the entire 3 mm window using a wide-field fluorescence macroscope (Axio Zoom.V16 Zeiss/Caenotec-Prof. Ralf Schnabel). The mouse was positioned in the center of the LCD screen. Images were captured at 20 Hz by a high-speed sCMOS camera (pco.edge 5.5) and recorded using the Encephalos software package (Caenotec-Prof. Ralf Schnabel).

All visual stimuli used for wide-field imaging were created and presented using COGENT (developed by John Romaya at the LON at the Wellcome Department of Imaging Neuroscience) running in MATLAB as described before^75^. In short, mice were positioned 14 cm away from a 122 × 68 cm LCD screen (Iiyama LE5564S-B1), resulting in a field-of-view of 144° x 86°. The screen had a resolution of 1920 × 1280 pixels running at refresh rate of 60 Hz. Stimuli used for pRF mapping were as described before^75^ and consisted of eccentricity-corrected angled bars (at 0°, 45°, 90° or 135°, 10° in diameter) of a checkerboard pattern on a gray (20 cd/m^2^) background. The total number of 58 stimuli were presented 10 times each for 0.5 s followed by a gray (20 cd/m^2^) ITI of 3.6 s. Natural stimuli used for wide-field occlusion experiments were similar as for two-photon experiments (described above). Both non-occluded and occluded images were presented 20 times for 1 s with an ITI of 4-6 s.

### Behavioral training and analysis

For behavioral training in the reward-based paradigm, water-deprived mice were trained to detect 4 out of 6 non-occluded images to make them familiar. The remaining 2 novel/untrained images were not shown during training. Mice were free to run on the wheel while images were presented for 2 s with a random inter-trial interval of 7-10 s. In the initial passive stage of the training, mice were rewarded with a 5 µl water drop 1 s after stimulus offset. When mice started to lick before the reward delivery, training became active, and a water reward was delivered only when the mice licked in the time window 1-2 s after stimulus offset. Imaging was performed when mice consistently managed to obtain a reward on >70% of trials (generally between 20-30 sessions of 100-200 trials each, ∼3750 trials). Mice continued to be trained in between imaging sessions until the experiments were completed.

The hit rate was defined as:

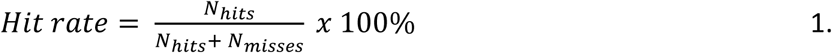

For behavioral training in the passive paradigm, mice were presented with 4 out of the 6 non-occluded images. The remaining 2 novel/untrained images were not shown during training. No lickspout was present and no rewards were delivered. Each mouse went through a 3-day training regime with 2 training sessions per day (morning and afternoon). Stimuli were presented for 1 s each with a random ITI of 4-6 s. Mice received a total of 3000 presentations per stimulus.

### Histology and confocal microscopy

To examine GCaMP expression, mice were perfused with 15 ml of ice-cold PBS followed by 30 ml of 4% paraformaldehyde (PFA). The brains were extracted and post-fixed for 2 h in 4% PFA at 4°C. Brains were cut in 50 μm slices and imaged on an Axioscan.Z1 slide scanner (ZEISS, Germany) at 20X (0.63 NA).

### Wide-field imaging analysis

Captured images (1600×1600 pixels, 20 Hz, 50 ms exposure) were down sampled to 800×800 pixels. Images were then smoothed using a Gaussian filter with a standard deviation of 2 pixels and stored in a trial based format (400 x 400 x time x number of trials). Analysis of pRF mapping and field-sign was performed as described in detail before^75^.

For analysis of occlusion experiments, images were semi-automatically registered using intensity-based rotation and translation to the pRF mapping session^75^. Using the pRF map, we extracted signals for the retinotopic region of V1 corresponding to the occluded quadrant of the screen (with a margin of 7 visual degrees to limit contamination from nearby non-occluded regions of V1). We averaged ΔF/F signals for all pixels in the occluded region and calculated the baseline-corrected (average activity in the 0.2 s before stimulus onset) mean traces across trials and conditions.

### Two-photon imaging analysis

We used the SpecSeg toolbox for preprocessing as described in detail before^76^. In short, we performed rigid motion correction using NoRMCorre^77^ followed by automated region-of-interest (ROI) selection based on cross-spectral power across pixels. After manual refinement, raw ROI signals were extracted and corrected for neuropil by subtracting the average pixel values in an area surrounding each ROI multiplied by 0.7. ΔF/F traces were made by subtracting a moving baseline (10th percentile over 5000 frame windows) and dividing by a linear fit over that baseline.

For RF analysis, we performed reverse correlation on the responses to the sparse noise stimuli to create a 2D response matrix for both the ON (white squares) and OFF (black squares) stimuli. We then fit a circular 2D Gaussian to the response matrix to estimate the RF center and size (full width at half maximum). The quality of the fit was estimated using r^2^ and the Signal-to-Noise Ration (SNR).

The SNR was defined as:

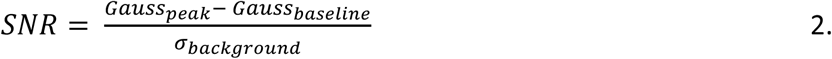

Where *Gauss_peak_* is the maximum of the Gaussian fit, *Gauss_baseline_* is the offset parameter and *σ_backbbround_* is the standard deviation of the background data points (>2 standard deviations away from the center of the Gaussian).

We included neurons if either their ON or their OFF RF met the following criteria: (i) their RF was reliable (r^2^ > 0.33 and SNR > 4) and (ii) the edge (based on the FWHM) of their RF was at least 2 visual degrees away from the bottom and right-side edges of the occluder (Fig. S2). For display purposes (Fig. 2 & S2), we used the ON RF for each neuron whose RF met criteria for both the ON and OFF RF map. For neurons recorded in naive and expert mice, we used RFs obtained from sparse noise stimuli shown directly before the occlusion stimuli. For the task data, we used RFs obtained from the expert RF mapping session by matching neurons between the expert and task dataset. Based on these criteria, 45% of all ROIs were included.

For 2P response strength analyses, we first z-scored ΔF/F signals for each neuron. To compare mean response traces for individual neurons to occlusion stimuli (Fig. 2C), we then averaged across repetitions per image followed by stimulus type (NO and O). To compute single cell average response strengths, we took the baseline-corrected (average 1 s before stimulus onset) mean response between t = 0.2-1 s after stimulus onset.

Average responses to surround suppression and annulus stimuli were computed as described above. We then normalized tuning curves for each neuron to their maximum responses and removed neurons with extreme minimum response values (<-20). We calculated the surround suppression index (SSI) as follows:

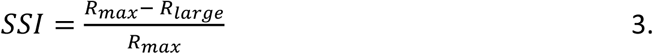

Where *R_max_* is the maximum response to any of the sizes and *R_larbbe_* is the response to the largest presented size. We calculated the annulus index (AI) as follows:

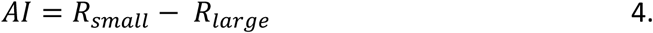

To estimate the relationship between population responses to non-occluded and occluded stimuli across different conditions (Naive, Expert, Task), we calculated the correlation coefficient (R) and the separation index (SI) between non-occluded and occluded responses. For R, we first computed single cell responses to non-occluded and occluded stimuli averaged across images. We then calculated the correlation coefficient for the entire population of neurons by correlating the non-occluded and occluded responses (x-axis and y-axis respectively in Fig. 2F). We calculated the SI as follows. First, we divided the 2D scatter plot (Fig. 2F) into four quadrants (2 x 2). We independently defined the borders between the quadrants for the x-axis and y-axis as the mean + 0.75*STD for the non-occluded and occluded responses, respectively. Second, we counted the number of neurons whose response profile was located in the top right quadrant. To estimate the degree to which non-occluded and occluded responses were separated, we compared the neuron count in the upper right quadrant of the real distribution to the same count in a non-correlated distribution with independent sampling for the x and y-coordinate. For this, we ran 10,000 repetitions of the count while each time randomly shuffling the non-occluded and occluded responses (creating random pairs of x and y values), resulting in a distribution of permutation counts. We calculated the separation index (SI) as follows:

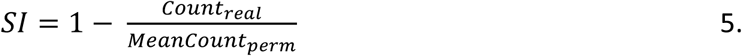

Where *Count_real_* is the upper right quadrant count from the real data and *MeanCount_perm_* is the distribution average of upper right quadrant counts from the permutation data. Values <-1 were clipped at –1. An SI value of –1 represents a separation lower than the chance distribution (i.e. anti-separation or correlation), 0 represents a lack of separation and 1 represents a separation higher than the chance distribution.

To correlate the strength of the response to the occluded images with the distance of RF edges to the occluder edge for each neuron, we took either the ON or the OFF RF of the neuron depending on which matched the criteria described above. If both the ON and OFF RFs matched the criteria, we took the ON RF. The distance of the RF edge to the occluder was determined as the shortest of the two possible occluder edges (horizontal or vertical).

We calculated response onset latency for each neuron as described before^32,78^ by fitting the sum of an exponentially modulated Gaussian and cumulative Gaussian to the activity time courses for non-occluded and occluded conditions during the task. We defined the onset latency as the time point at which the fitted curve reached 33% of its maximum. We only included neurons with a high-quality fit (r^2^ > 0.5) and an average response (t = 0.2-1 s) greater than 0.

To estimate the contribution of locomotion to responses and plasticity we used two independent procedures. First, we performed linear regression for each neuron by using locomotion speed as a covariate and neuronal activity as a response variable. We then used the residuals of the model for all subsequent analyses. Second, we repeated the analyses after removing all trials in which the mouse had an average running speed > 2 cm/s during the stimulus window (t = 0.2-1 s).

We defined stimulus selectivity across the 4 different natural images by calculating lifetime sparseness for each neuron as follows:

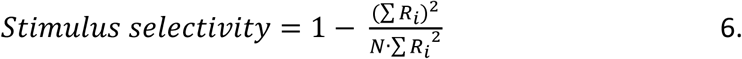

Where *Ri* represents the neuron’s response to the *i*-th stimulus, and *N* is the total number of stimuli. We only included responsive neurons (response > 0.3 for that condition).

We calculated selectivity for novel stimuli compared to familiar stimuli in neurons that are responsive to novel stimuli (response > 0.5). For each neuron, we used responses to image pairs in which one image was novel and the other was familiar. For each image pair, we computed a selectivity index as follows:

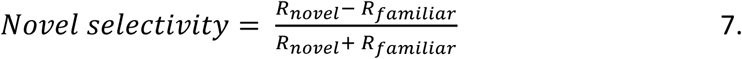

where *R_novel_* and *R_familiar_* represent the neuron’s response to the novel and familiar image in the pair, respectively. Only pairs for which both responses were greater than zero were included.

We matched neurons across chronically recorded sessions using the chronic matching module of SpecSeg^76^. To correlate the change in response strength between naive and expert conditions to the selectivity to the non-occluded stimulus for individual neurons, we computed the correlation coefficient using the population of neurons that were found back in both sessions.

We calculated the activity ratio based on reverse correlations (Fig. S12) as follows. For each neuron, we computed the reverse correlation across 3,600 natural images, as the weighted sum of the response to each image (averaged in the time window 0-500ms after the stimulus onset, and zero-centered) and the image structure computed using Shannon’s entropy. Image structure, i.e. the entropy, was computed locally in bins of non-overlapping 30×30 pixels, using *skimage*^79^. This resulted in a single image for each neuron, corresponding to their profile of spatial activation. Then, population images were obtained by averaging the individual images of all neurons, separately for each condition (Fig. S12A). Finally, for the population images (Fig. S12B) we computed the ratio of activity elicited by image parts outside the classical RF (cRF) and the parts inside the cRF. The cRF region was defined at the population level, by summing up the RFs of all the neurons, thresholding the obtained result at the 95^th^ percentile. The SEM of the ratios was computed by bootstrap resampling (1,000 iterations), while statistical differences were tested for significance using a permutation test (1,000 iterations) and corrected for multiple comparisons with Bonferroni.

### Decoding analysis

For population decoding of mouse 2P data and monkey MUA data, we performed LDA using the MATLAB function fitcdiscr (default parameters) on baseline-corrected neuronal responses measured between 0.2 and 1 s after natural image onset. For decoding over time using mouse 2P data, the same procedure was applied using the baseline-corrected ΔF/F value from each imaging frame. Decoding over time for monkey MUA data was adapted from Papale et al. (2023)^27^. Decoding of human fMRI data was adapted from Smith & Muckli (2010)^25^. To control for locomotion-related activity, the influence of running speed was regressed from neuronal responses prior to decoding. On each decoding run, 300 neurons were randomly selected, corresponding to the smallest neuronal population size available across experimental groups. Trials were then divided separately for each image into training and test datasets (50%/50%), ensuring balanced representation of all stimulus identities in both datasets. To prevent train-test leakage, z-scoring parameters were estimated from the training dataset only and subsequently applied to both the training and test datasets. We then trained an LDA decoder on the training dataset and tested the decoder on the remaining trials. We repeated this procedure 200 times to account for variability between trials and neuron selection. We built two classifiers, one on non-occluded trials (NO model) and one on occluded trials (O model). We then tested each classifier on both conditions to yield four accuracy scores.

Statistical significance of decoding performance was assessed using permutation testing as described before^27^. For each decoding condition, a null distribution was generated by repeating the complete decoding procedure 1000 times with shuffled stimulus labels. To improve estimation of extreme tail probabilities, a generalized Pareto distribution was fitted to the upper 30% of the null distribution^80^. P-values were calculated from the position of the observed decoding accuracy relative to the fitted null distribution.

### Statistical analyses

All statistical details for each experiment are shown in figures and figure legends and in the statistical table. Unless specified otherwise, statistical analyses were performed using LMEM (‘fitlme’ function in MATLAB). For LMEM, we considered the response parameter (e.g. visual response magnitude) as the dependent variable, mouse as random effect and the condition (e.g. Naive vs Expert) as the fixed effect. LMEM accounts for the hierarchical correlation structure in the data (i.e. neurons and mice). When the model resulted in a Hessian error (i.e. random effect could not be estimated), we iteratively removed outlier neurons (i.e. neurons with residuals <-2.5 or >2.5 z-score) before refitting the model. We performed a one-way ANOVA on the LMEM followed by a post hoc coefTest using Tukey’s HSD to correct for multiple comparisons.

### Computational model

#### Circuit model

We modeled the opposing plasticity in PyCs using a simplified rate-based circuit model outlined in Figure 7A, consisting of a two-compartment PyC *y* and a single-compartment PV interneuron *p* providing feedforward divisive inhibition^47^. In this model, the apical and basal PyC compartments are functional rather than biophysical dynamical compartments, and each model instance represents the mean response dynamics of a phenomenological PyC–PV circuit corresponding to one experimentally observed average response-transition class, rather than an individual biophysical neuron.

Firing rate of the PyC *y* was modeled as a nonlinear leaky integrator of the apically-modulated somatic activation *γγb*, apical drive *β*, and an independent baseline current 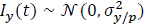 randomly sampled at each timestep. The PyC firing rate was then passed through a rectified linear activation function, *φ*(*u*) = max{0, *u*}.

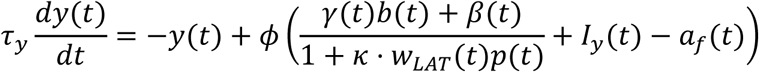

The basal compartment 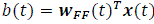 of the PyC received feedforward inputs *x*(*t*) ∈ {0,1}^3^ via dedicated basal feedforward synapses ***w****_FF_*(*t*) ∈ (0,1)^3^ for feedforward inputs corresponding to each of the experimental stimuli. The integrated feedforward and contextual inputs were normalized by divisive feedforward inhibition^47^ from the PV cell *p* via an inhibitory somatic synapse *w_LAT_*(*t*) ∈ [0,1], with parameter *κ* setting the maximum possible scale of divisive inhibition. The PyC also accumulated adaptation currents *a* with persistent stimulation. Biologically realistic short-term (within-trial) adaptation of neural responses was a result of the fast adaptation current *a_f_*, acting as an additional leak term on the PyC dynamics. Furthermore, the effective rate of feedforward plasticity was directly proportional to a slow adaptation current *a_s_* that accumulated basal activity *b*(*t*) over several trials, *τ_as_* >> *τ_af_* >> *τ_y_*.

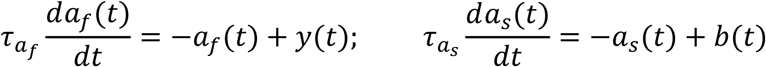

The apical dendritic compartment *d* of the PyC received generalized contextual inputs *c* ∈ (0,1)^3^, with 60% overlap in terms of the individual apical feedback synapses ***w****_FB_* ∈ (0,1)^3^ stimulated by different contexts corresponding to individual experimental stimuli, *d*(*t*) = ***w****_FB_*(*t*)*^T^c*(*t*). To account for nonlinear nature of apical feedback which can amplify somatic activity^81^, but also drive somatic firing when feedback is sufficiently strong^63–65^, we treated the apical multiplicative gain *γ* and apical drive *β* as separate terms.

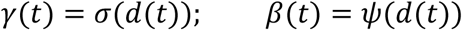

Apical gain *γ* was modeled with a thresholded, shifted sigmoid function 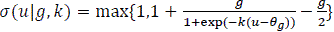, to ensure that apical feedback could enhance, but not suppress somatic activity, with parameters *g* and *k* controlling the apical gain ceiling, and steepness of the gain function. Apical feedback acted as multiplicative gain on somatic activity as soon as apical activation crossed a threshold *θ_g_* shared across cells. Apical drive *β* was modeled with a shifted rectified linear function *ψ*(*u*|*θ_d_*) = max{0, *u* − *θ_d_*}, only contributing to neuronal activity once apical activation crossed a threshold *θ_d_*. The apical activation necessary for gain modulation of somatic activity was always lower than the apical activation necessary for apical drive *θ_bb_* < *θ_d_*.

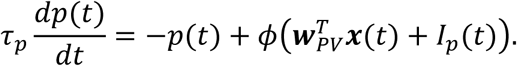

We modeled feedforward divisive inhibition, with a point PV interneuron *p* receiving the same feedforward input *x* as the pyramidal cell, with distinct synapses for each of the feedforward inputs ***w****_PV_* ∈ (0,1)^3^. The PV cell also received a small random baseline current 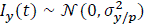 sampled independently at each timestep. The firing rate of the PV *p* was also modeled as a nonlinear leaky integrator, with the same nonlinearity as PyC. The firing rates of both PyC and PV were clipped to 1.0, representing the maximum possible firing rate for each cell type.

#### Plasticity rules

In our model, PyC synapses undergo plasticity according to three different learning rules highlighted by the different colors in Figure 7A, excitatory inputs onto PV neurons do not change with experience. Updates for all potentiating weights are softly bounded by a factor of (1 − *w*), and by a factor of *w* for adapting weights, such that synaptic weights range between 0-1.

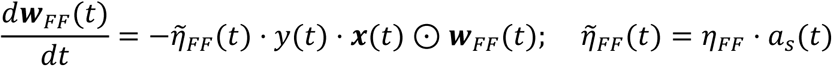

The PyC feedforward synapses ***w****_FF_* undergo selective soft-bounded depression as function of postsynaptic activity *y*, at a dynamic rate 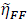. Based on the increased selectivity of NO responses to both familiar and novel images after familiarization (Fig. 2H, 4H), we hypothesized that the effective learning rate 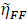 scales with breadth of naive feedforward tuning of PyCs. In our model broadly tuned PyCs depress more rapidly than narrowly tuned PyCs. This phenomenon emerges due to the slow timescale *τ_as_* which makes *a_s_* accumulate over several independent stimulus presentations. Through dynamic direct modulation of *η_FF_*, higher *a_s_* accumulated over several trials in frequently active broadly tuned cells results in a consistently higher 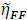, compared to less frequently active narrowly tuned cells whole *a_s_*, and consequently 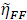 is lower on average across time.

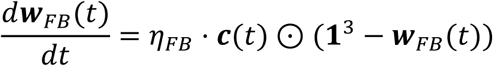

Conversely, PyC apical feedback synapses ***ww****_FB_* strengthen proportionally to generalized presynaptic contextual input c, at a rate *η_FB_*. Because contextual signals overlap in the set of apical synapses they stimulate, this simple rule enables generalized strengthening of contextual responses to novel stimuli.

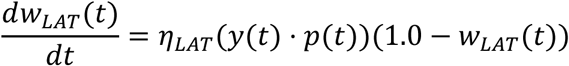

Finally, the weight of the inhibitory synapse *w_LAT_* onto the PyC evolves according to a soft-bounded Hebbian plasticity rule, with learning rate *η_LAT_*. This rule contributes to long-term experience-dependent adaptation of non-occluded responses to novel stimuli.

#### Simulation details

We studied PyC responses to two familiar inputs (*E*_1_, *E*_2_) and one novel input (*E*_3_). Each input matrix representing non-occluded (NO) inputs 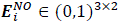 was constructed by stacking a one-hot vector 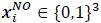 representing image-specific feedforward inputs, and a dense vector *c_i_* ∈ (0,1)^3^ representing generalizing contextual inputs stimulating all apical synapses simultaneously. Specifically, the contextual information for each image was simulated with a permutation of the contextual vector [1.0,0.6,0.6]/2.2 to model broader generalized nature of ecRF feedback compared to cRF feedforward inputs. Occluded images 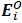 were modeled by replacing the image-specific feedforward input with a vector of zeros, 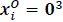. We simulated our model in discrete time, yielding *T* frames per trial, consisting of 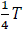 pre-stimulus ITI frames, 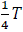 stimulus presentation frames, and 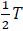 post-ITI frames. During the ITI, the model input consisted of only zeros *E^ITI^* = **0**^3×2^, representing absence of visual input.

We followed the same experimental protocol as the real experiment (Figure 7A), first testing responses of naive neurons to both NO and O versions of all inputs 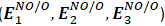. Subsequently, we familiarized neurons with the NO versions of the familiar images only, by showing *N_train_* presentations of each of the familiar images 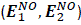 in alternating order. After familiarization, we tested the responses of expert neurons to all inputs again 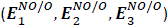. Plasticity rules were only activated during the familiarization phase—in both naive and expert testing sessions, PyC synaptic weights (***w****_FF_*, ***w****_FB_*, *w_LAT_*) were frozen to isolate the effect of plasticity during the familiarization phase. In the expert testing session we included three additional perturbation conditions during which we tested the neuronal response to all inputs again—silencing PV by clamping *p*(*t*) to 0; silencing the contextual feedback inputs by clamping contextual input vector to 0^3^; and silencing both PV and contextual feedback simultaneously. These perturbations allowed us to better understand the role of different components of the model, and generate hypotheses for future experiments which could perform these perturbations using *in vivo* optogenetics.

We modeled the three most common response transitions observed *in vivo*, separately for familiar and novel images, yielding six separate mean-field transition models. –NO transition was defined as a decrease in NO response greater than the change in O response; +O transition as an increase in O response greater than the change in NO response, and +NO transition as an increase in NO response greater than the change in O response. Through a combination of constrained random parameter search, and hypothesis-driven manual fine-tuning, we defined a set of simulation parameters shared across all transition models (Table S1), as well as, a second smaller set of PyC and PV parameters defining each mean-field transition (Table S2).

Each mean field firing rate was standardized based on the mean and standard deviation of the neuron’s firing rate across all the pre-stimulus periods in the naive testing phase.

We ran our simulations in discrete time, using Euler integration, yielding a discrete-time leaky integration version of our differential equations for the PyC firing rate *y*, PV firing rate *p*, and adaptation current *a*.

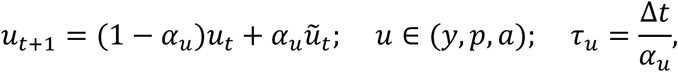

where 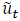 is the instantaneous activation. Here the leak-rate is determined by the constants *α_u_*, which are related to the continuous time-constants *τ_u_* via the time-discretization Δ*t*.

#### Robustness analysis

To verify the robustness of our model to the exact parameter values and ratios between parameters, we conducted a robustness analysis, where we varied one continuous parameter at a time around each of the six mean-field models’ parameter initializations. Concretely, every scalar model parameter was multiplied by factors ranging from 0.05x to 5x. The leak rates *α_y_* and *α_p_*, which are fractions, were capped at one. To check whether the model was sensitive to the magnitude of the fitted weights while preserving the fitted proportions, each complete set of weights (***ww****_FF_*, ***ww****_FB_*, ***ww****_PV_*, and *w_LAT_*) was scaled simultaneously by the same factors, and the scaled weights were clipped to the allowed weight interval [0,1] enforced throughout modeling. Furthermore, every individual [0-1] weight coordinate was also independently set to the absolute values between 0.01 and 1.0, spanning the full allowed weight range independently of each fitted weight’s initial magnitude. With each robustness check, all unperturbed parameters remained at their fitted mean-field transition values. This robustness validation design contained 1,965 perturbations across the six mean-field transition models.

To evaluate the outcomes of the robustness analysis, we measured the effects of each perturbation on 1) the magnitude of the perturbed transition vectors for each transition relative to the unperturbed case, and 2) whether the new perturbed transition vector ended up in the same transition quadrant as the unperturbed transition (+NO / –NO / +O). We considered a transition-parameter combination robust if the perturbed transition vector showed at least 25% magnitude of the original transition vector in the same transition sector as the original average transition.

This analysis shows that while partially arbitrary, the transitions we observe consistently arise because of our proposed plasticity rules acting on the canonical PyC-PV circuit, rather than because of the exact parameter setting (Fig. S14).

## Supplemental figures

**Figure S1.**
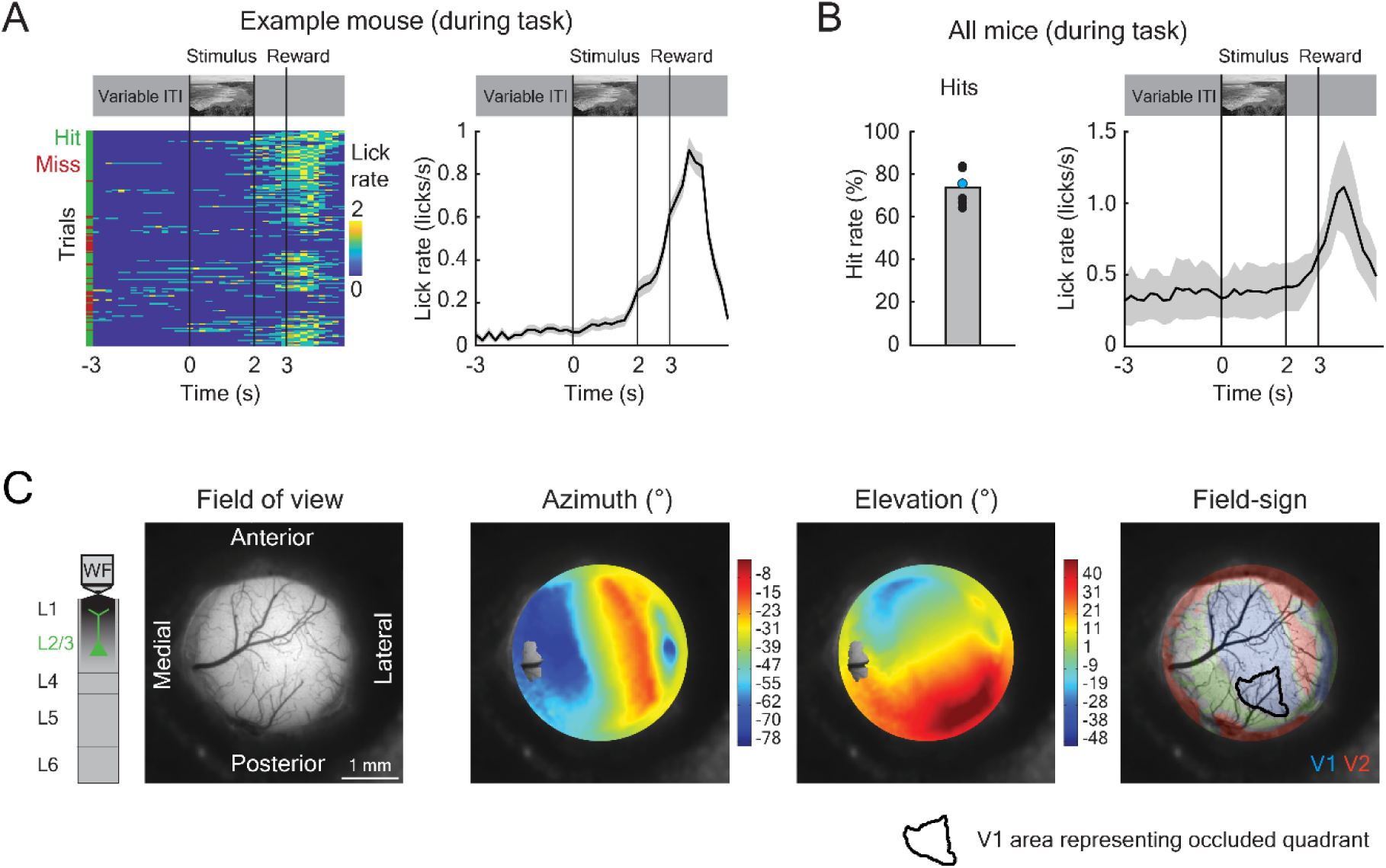
Task behavior and widefield population receptive field maps. A) Schematic of task structure and licking behavior for an example mouse (blue dot in B). Water restricted mice were required to lick in the interval of 1-2 s after disappearance of a non-occluded natural scene to obtain a water reward. Left: lick rate for individual trials. Right: trial-averaged average lick rate. B) Left: average hit rate for individual mice during the task session. The blue dots corresponds to the example mouse in A. Right: trial-averaged lick rate (mean ± SEM across 6 mice). C) Cortical wide-field maps for an example mouse showing from left to right: the imaging field of view, visual azimuth, visual elevation and visual field-sign overlaid on the brain imaged through the glass window. The large blue patch in the field-sign image represents V1 while the red patches represent higher visual areas (V2).

**Figure S2.**
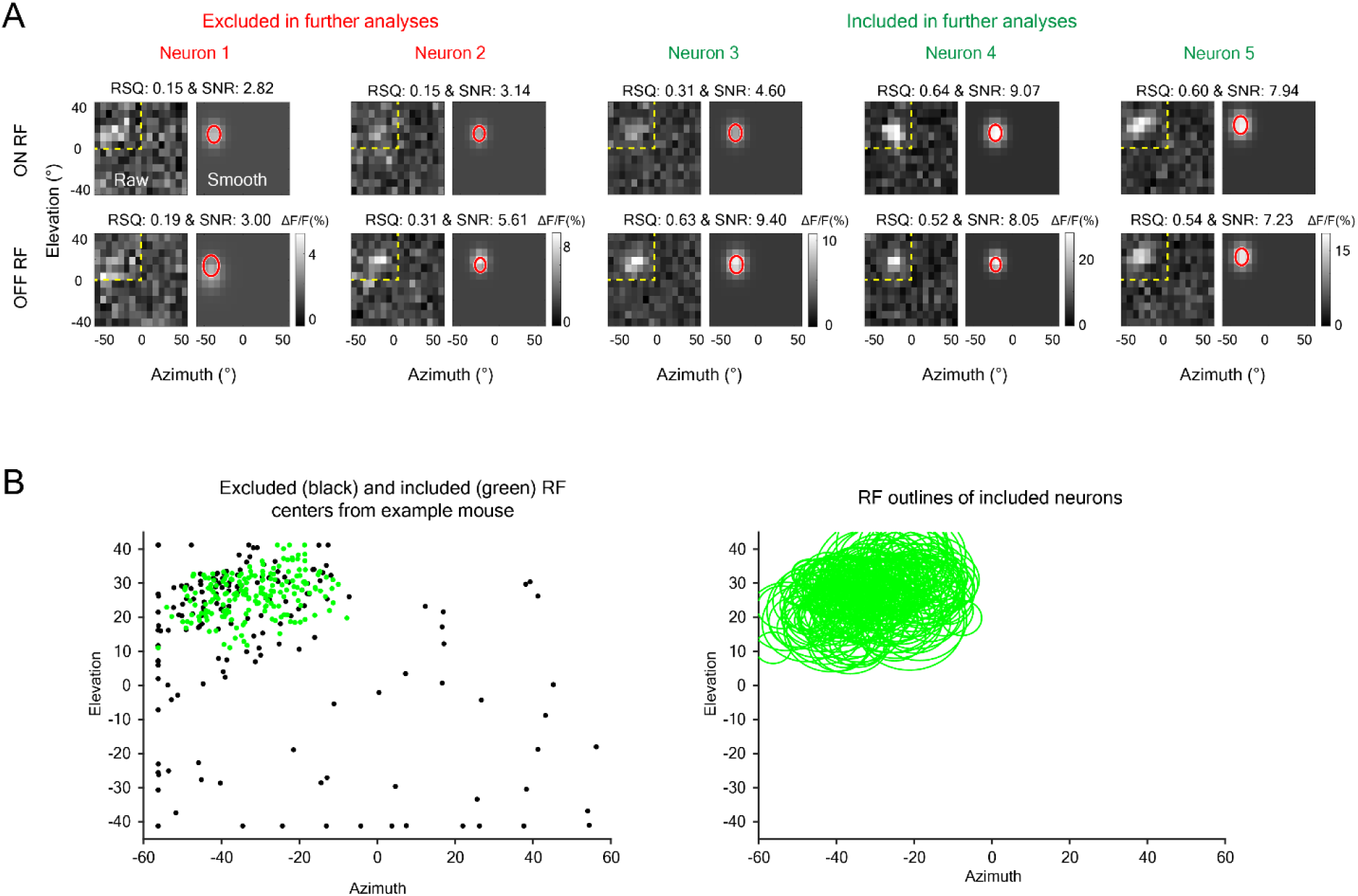
Receptive field inclusion procedure. A) Example receptive fields for neurons excluded (neurons 1 and 2, in red) or included (neurons 3-5, in green) in further analyses. B) Receptive field centers (left) and outlines (right) of neurons for one example mouse. Green and black dots respectively represent neurons included or not included in further analyses, based on quality or RF location.

**Figure S3.**
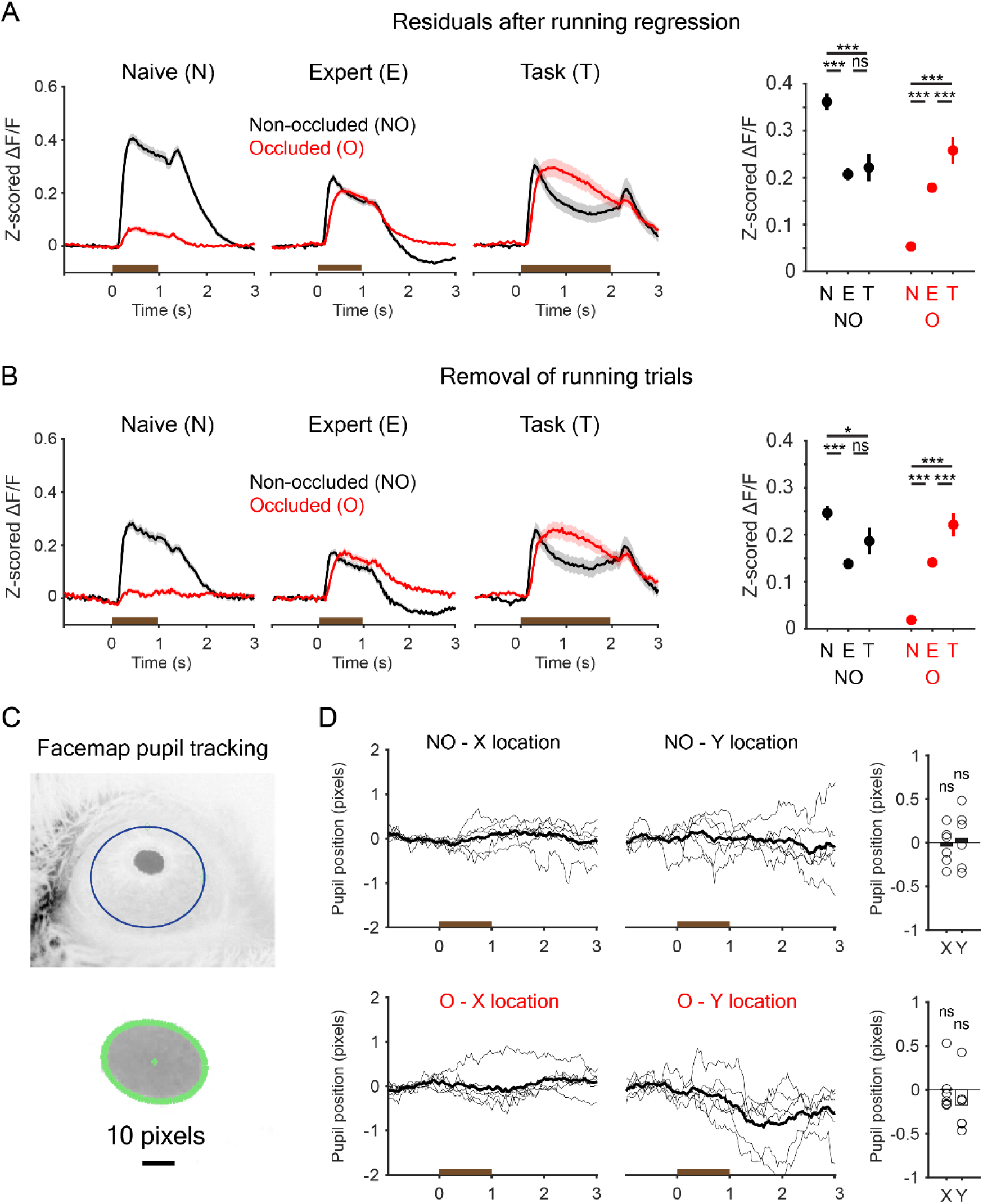
Locomotion and pupil position do not explain experience-dependent plasticity. A) Left: Schematic and z-scored ΔF/F traces averaged across neurons and stimuli for all conditions after regressing out locomotion speed and performing analyses on the residual data. Brown bars indicate the presence of the visual stimulus. Right: neuron-averaged response strength across stimuli and conditions (mean ± SEM, averaged between t = 0.2-1 s) for neurons after regressing out locomotion speed. N = 841 neurons (Naive), 844 neurons (Expert) and 300 neurons (Task) from 6 mice. LMEM with post hoc Tukey HSD for all comparisons here and below, ***: p<0.001, **: p<0.01, *: p<0.05, ns: not significant. B) As in A, but for data after removal of locomotion trials (average speed > 2 cm/s). C) Top: example image of pupil tracking using Facemap^82^. The blue circle represents the eye ROI drawn by the user within which Facemap will track the pupil. Bottom: zoomed in image of the pupil with the tracked pupil ROI (green circle) for an example frame. D) Traces for pupil position along the X (left) and Y (center) axes of the tracked pupil center for non-occluded (NO, top) and occluded (O, bottom) images. The brown bar indicates stimulus presence. Right panels show the average pupil position per mouse (black circles) during stimulus presentation (t = 0-1 s) relative to baseline (t < 0 s). Signrank test against zero, ns: not significant.

**Figure S4.**
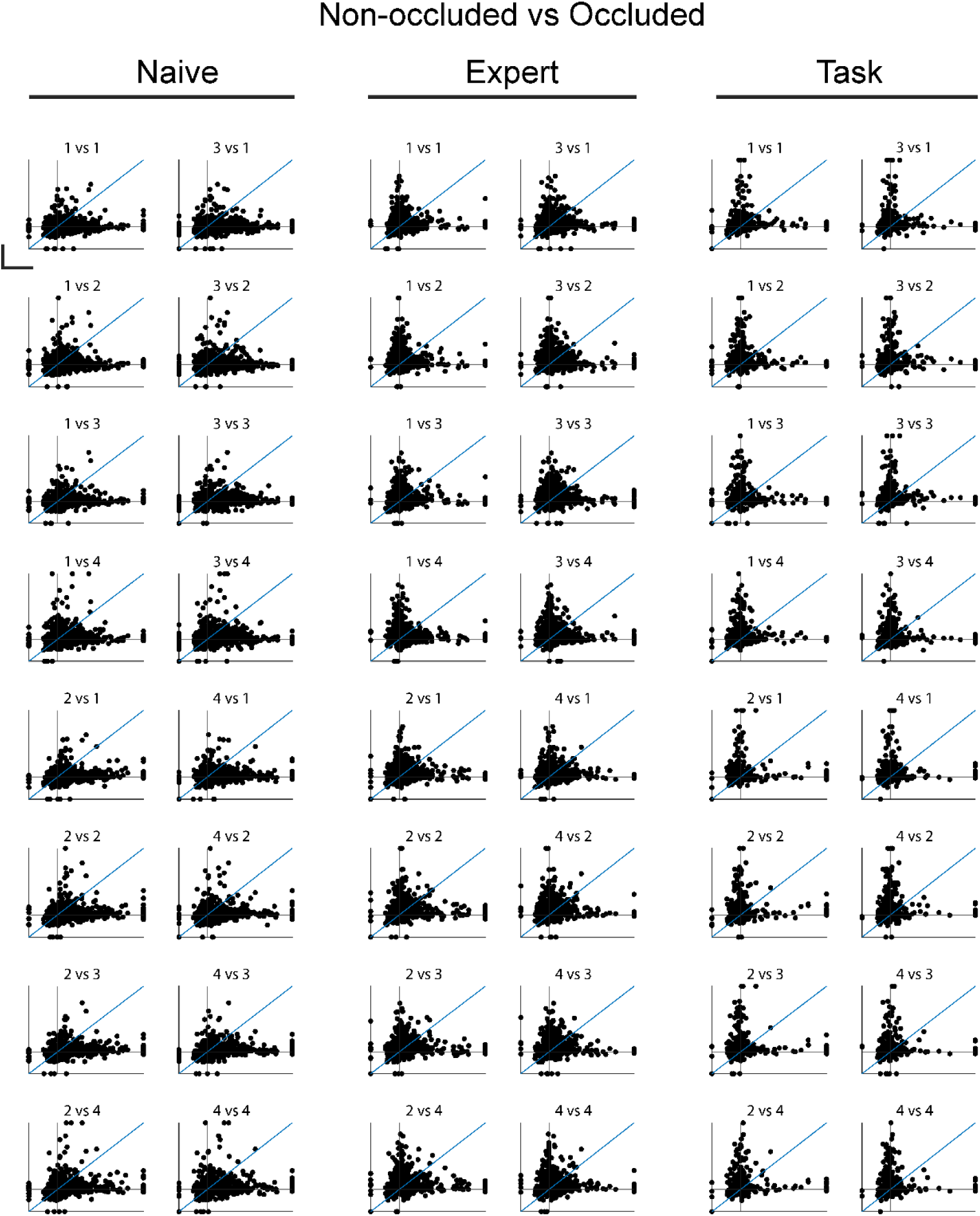
Relationship between responses to non-occluded and occluded images. Scatter plots showing the relationship between responses to pairs of non-occluded and occluded images. Panel titles indicate the image comparisons made (non-occluded on x-axis vs. occluded on y-axis). Axes are capped at –0.5 and 2.5 z-score. Scale bars, 1 z-scored ΔF/F.

**Figure S5.**
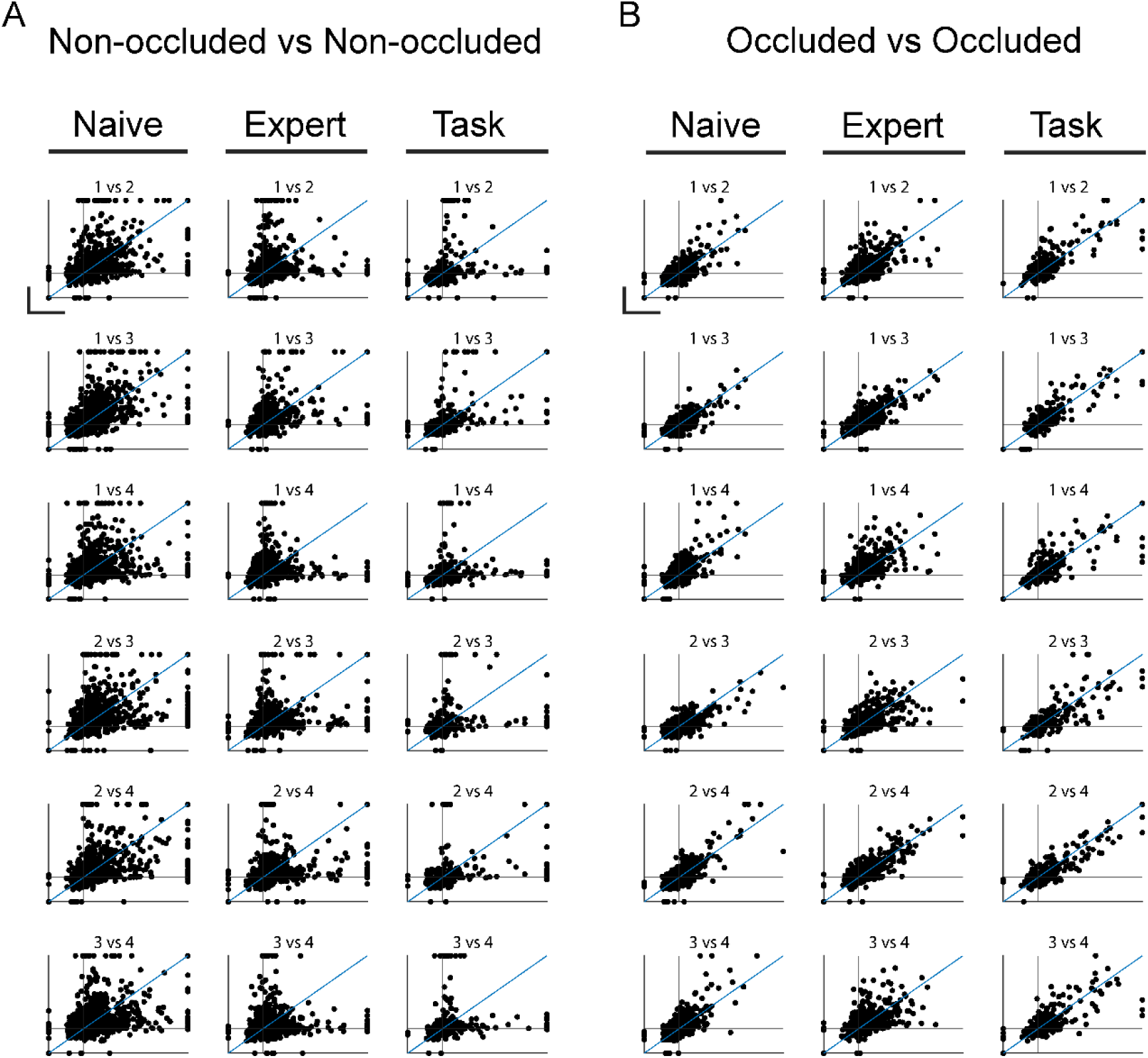
Relationship between responses to non-occluded and occluded images. A) Scatter plots showing the relationship between responses to pairs of non-occluded images. Panel titles indicate the image comparisons made (non-occluded vs non-occluded). Axes are capped at –0.5 and 2.5 z-score. Scale bars, 1 z-scored ΔF/F. B) As in A, but for pairs of occluded images.

**Figure S6.**
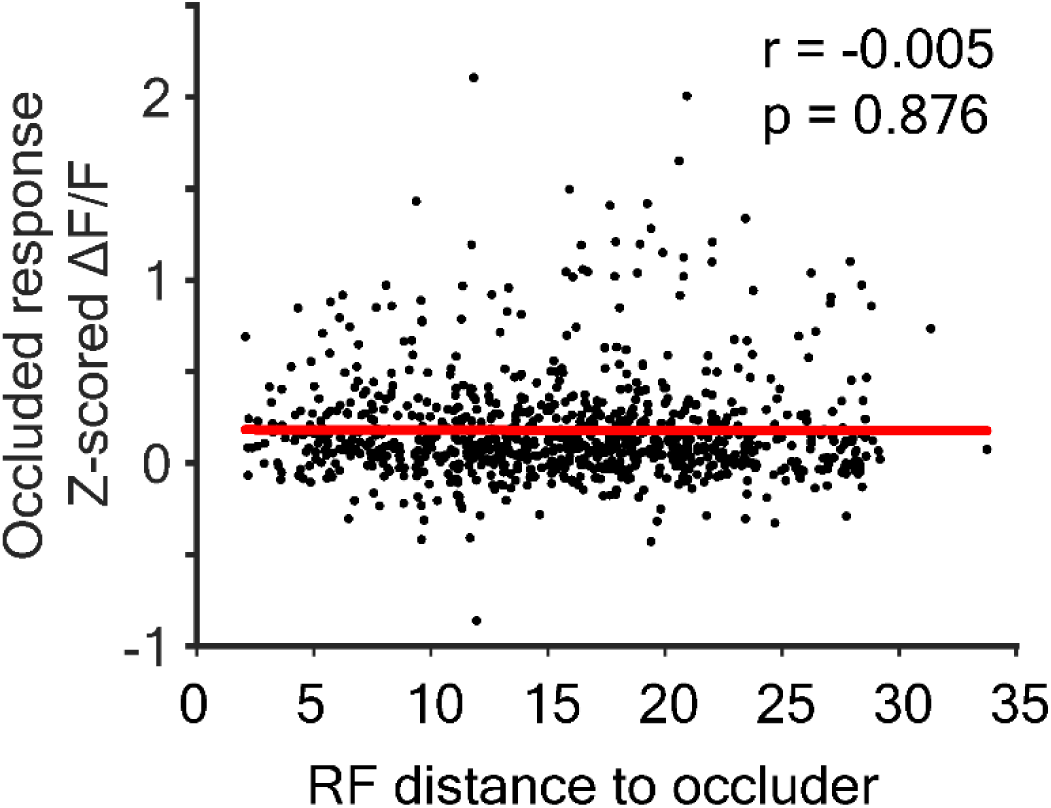
Relationship between occluded response strength and RF distance to the occlude in expert mice. Scatter plot showing the relationship between the distance between RF edge and the border of the occluder and the response to occluded stimuli. The linear regression line is indicated in red.

**Figure S7.**
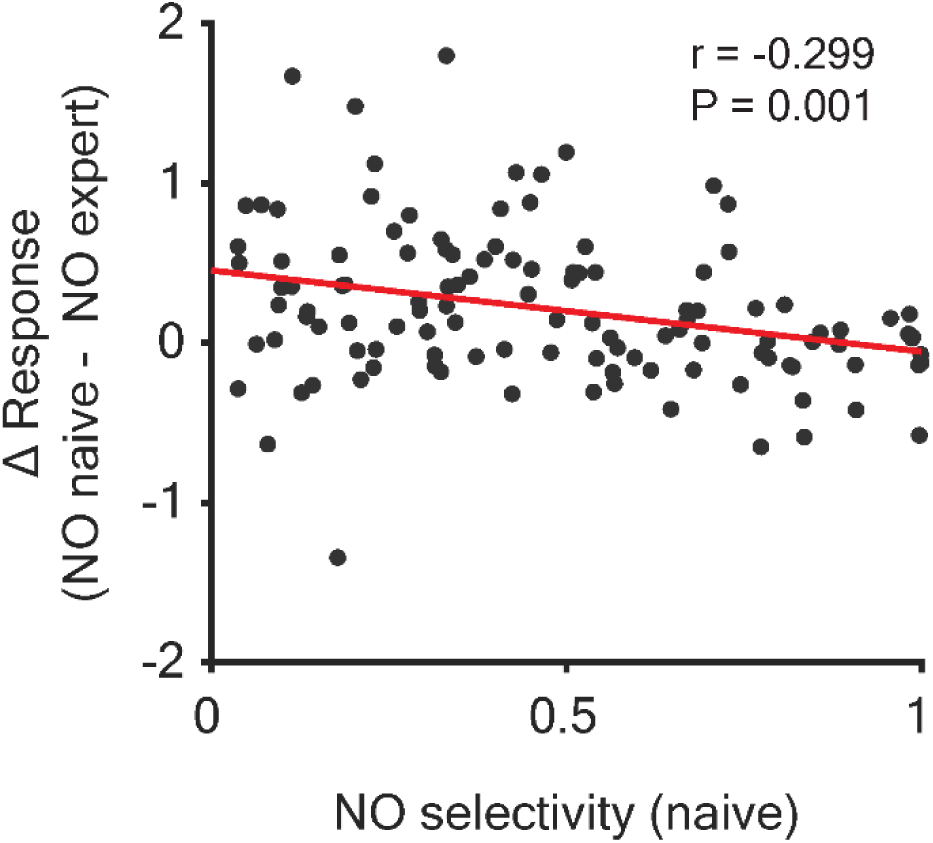
Relationship between non-occluded selectivity and plasticity. Scatter plot showing the relationship between selectivity to non-occluded images and the change in response strength from Naive to Expert conditions (n = 117 PyCs). NO, non-occluded.

**Figure S8.**
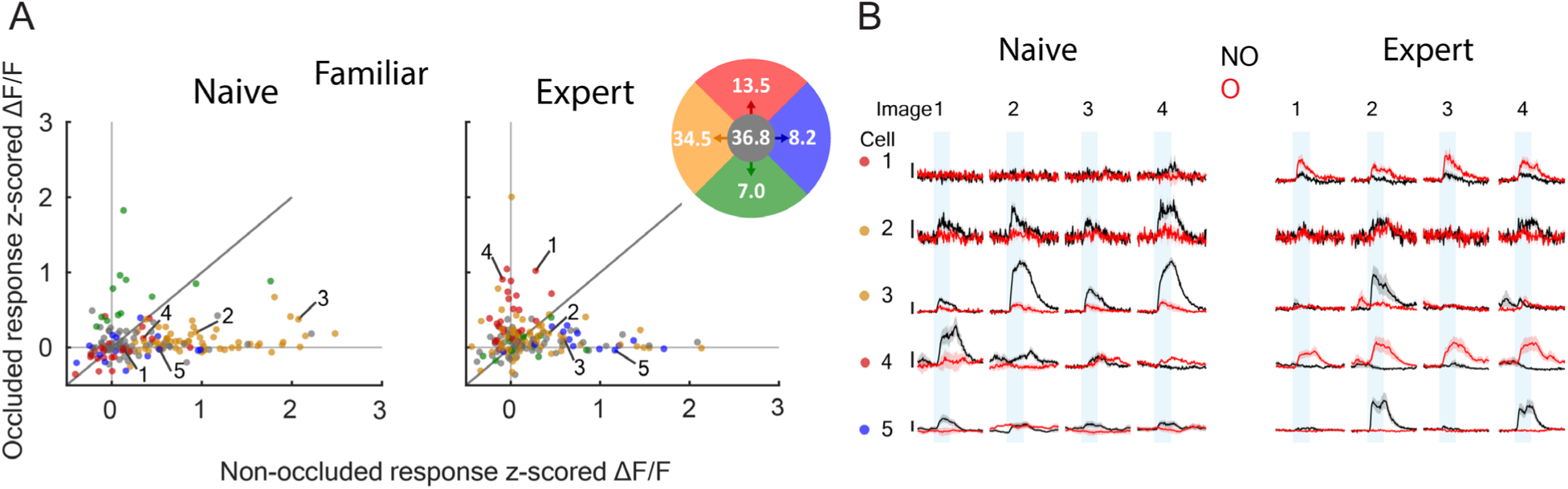
Neurons matched across chronic imaging sessions for Naive vs Expert NO-O responses. A) Scatter plot showing average z-scored ΔF/F responses (t = 0.2-1 s) to the non-occluded (x-axis) and occluded (y-axis) stimulus of chronically matched cells in naive and expert mice, color coded for the dominant direction in which their responses transitioned. N = 171 neurons from 6 mice. NO, non-occluded; O, occluded. The colored disc displays the percentages of the neurons transitioning in each direction. B) Neuronal responses of individual, chronically matched example neurons in naive (left) and expert (right) mice. NO, non-occluded; O, occluded. Scale bar,z 1 z-scored ΔF/F. Traces represent mean ± SEM over trials. The shaded bars indicate stimulus epochs.

**Fig. S9.**
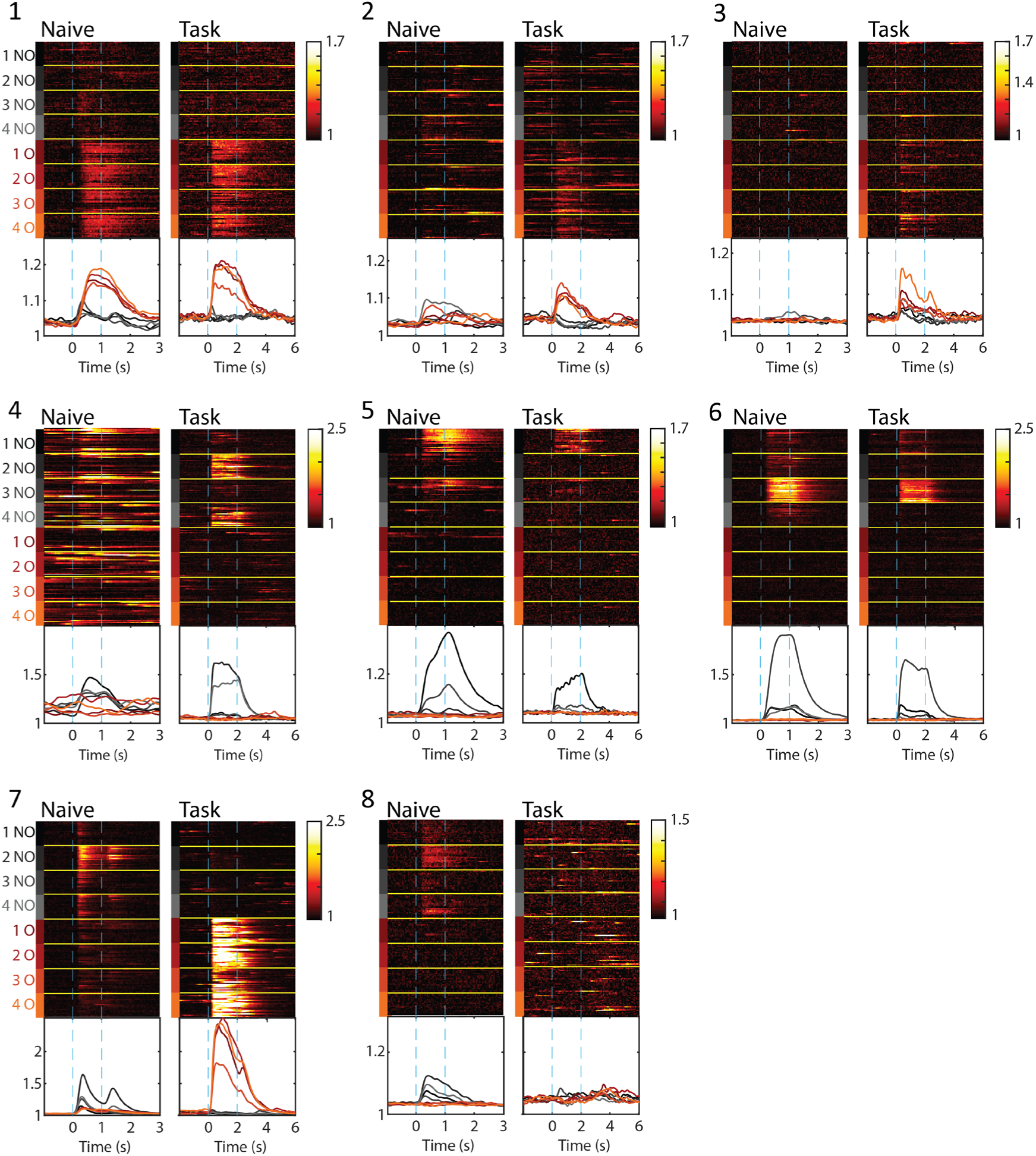
Per-trial responses of the chronic matched neurons from Fig. 2J. Each panel shows the activity (ΔF/F) of a single neuron across all trials, grouped by stimulus condition. *Upper section:* Individual trial responses sorted by stimulus type – non-occluded stimuli (NO 1–4, gray shades) and occluded stimuli (O 1–4, red shades). *Lower section:* Trial-averaged response for each stimulus condition.

**Figure S10.**
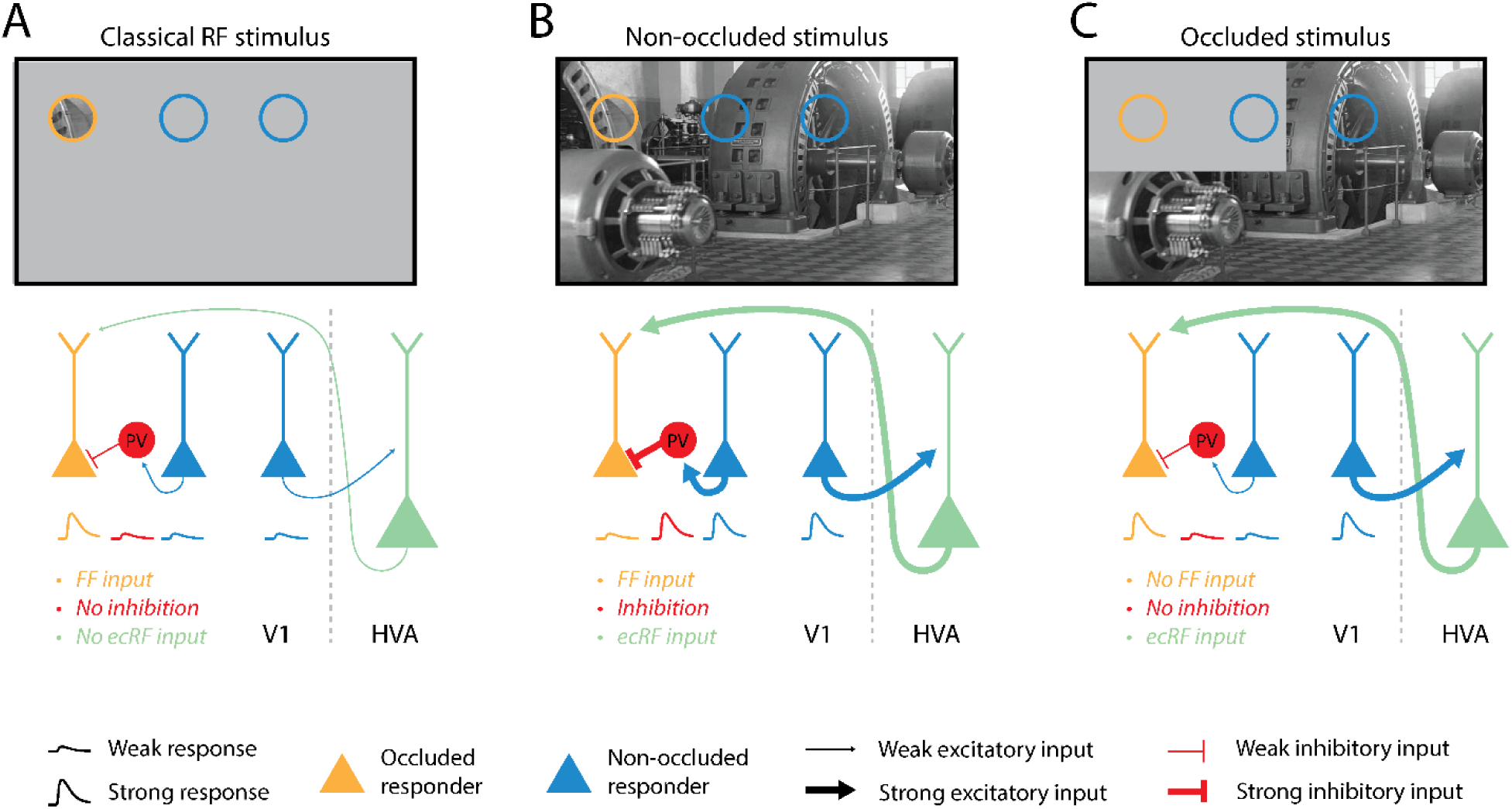
Mechanism underlying selective responses to occluded images. A) A small stimulus in the classical (cRF) drives the left PyC (orange, occluded responder). The PyC receives little inhibition (red) and no extra-classical RF (ecRF) inputs (green). B) A full visual scene gives input to the orange PyC, but also activates other PyCs (blue, non-occluded responders) that activate local interneurons (red), inhibiting the response of the orange PyC, despite ecRF input. C) An occluded visual scene, lacking of feedforward input, reduces local inhibition. Combined with ecRF inputs, the orange PyC is activated.

**Figure S11.**
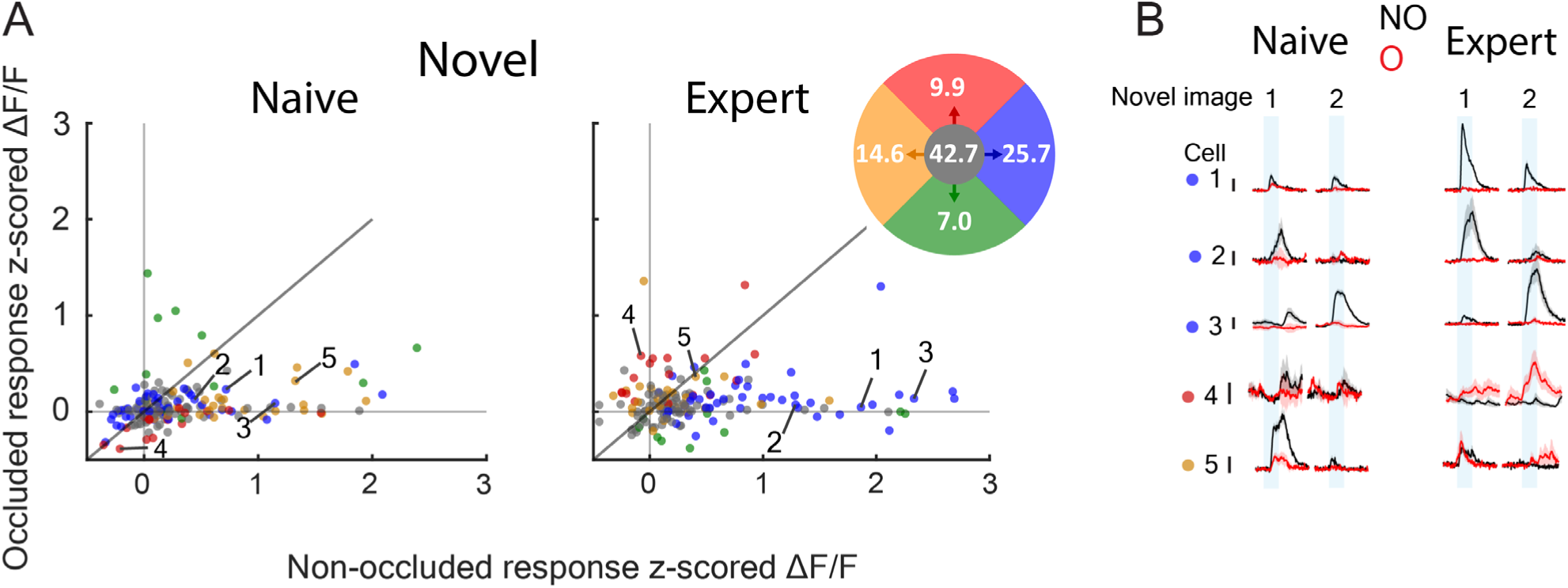
Neurons matched across chronic imaging sessions for Naive vs Expert NO-O responses. A) Scatter plot showing average z-scored ΔF/F responses (t = 0.2-1 s) to the non-occluded (x-axis) and occluded (y-axis) novel image stimulus of chronically matched cells in naive and expert mice, color coded for the dominant direction in which their responses transitioned. N = 171 neurons from 6 mice. NO, non-occluded; O, occluded. The colored disc displays the percentages of the neurons transitioning in each direction. B) Neuronal responses of individual, chronically matched example neurons in naive (left) and expert (right) mice. NO, non-occluded; O, occluded. Scale bar, 1 z-scored ΔF/F. Traces represent mean ± SEM over trials. The shaded bars indicate stimulus epochs.

**Figure S12.**
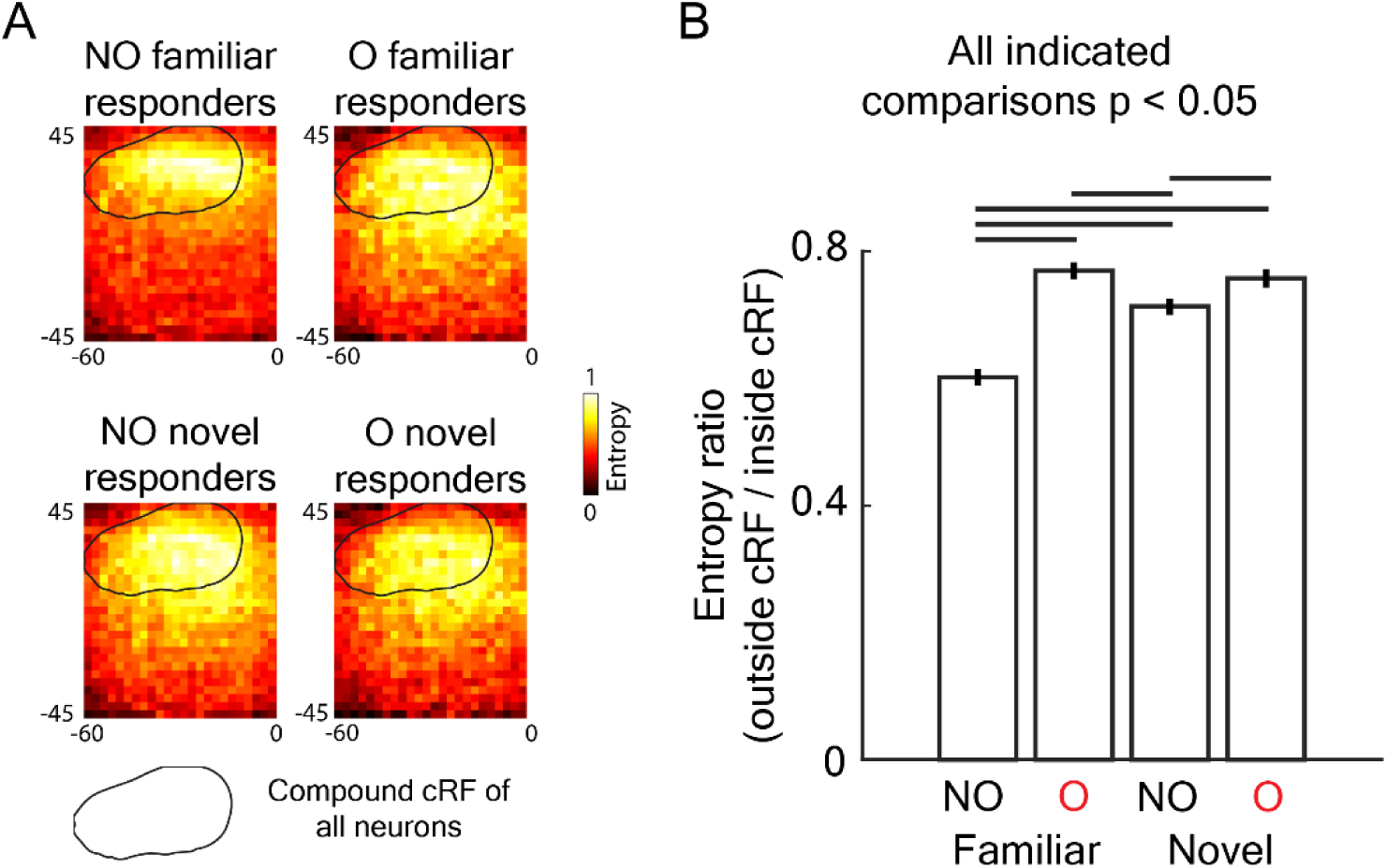
Image structure representations and outside-inside receptive field entropy ratios across L2/3 PyC categories. A) Average image structure representations per neuronal response type. Panels show averaged entropy across all reverse correlated (non-occluded) natural images, which captures the complexity of pixel intensities within an image region, for the indicated categories of PyCs (selected based on response > 0.5 for the indicated stimulus). The black outline represents the compound cRF of all neurons. B) Ratios of outside to inside entropy values per category. Values were computed as the ratio of entropy elicited by image parts outside the cRF and the parts inside the cRF. N = 277 neurons (NO familiar), 63 neurons (O familiar), 433 neurons (NO novel) and 52 neurons (O novel) from 7 mice. All indicated comparisons p < 0.05, permutation test with Bonferroni correction.

**Fig. S13.**
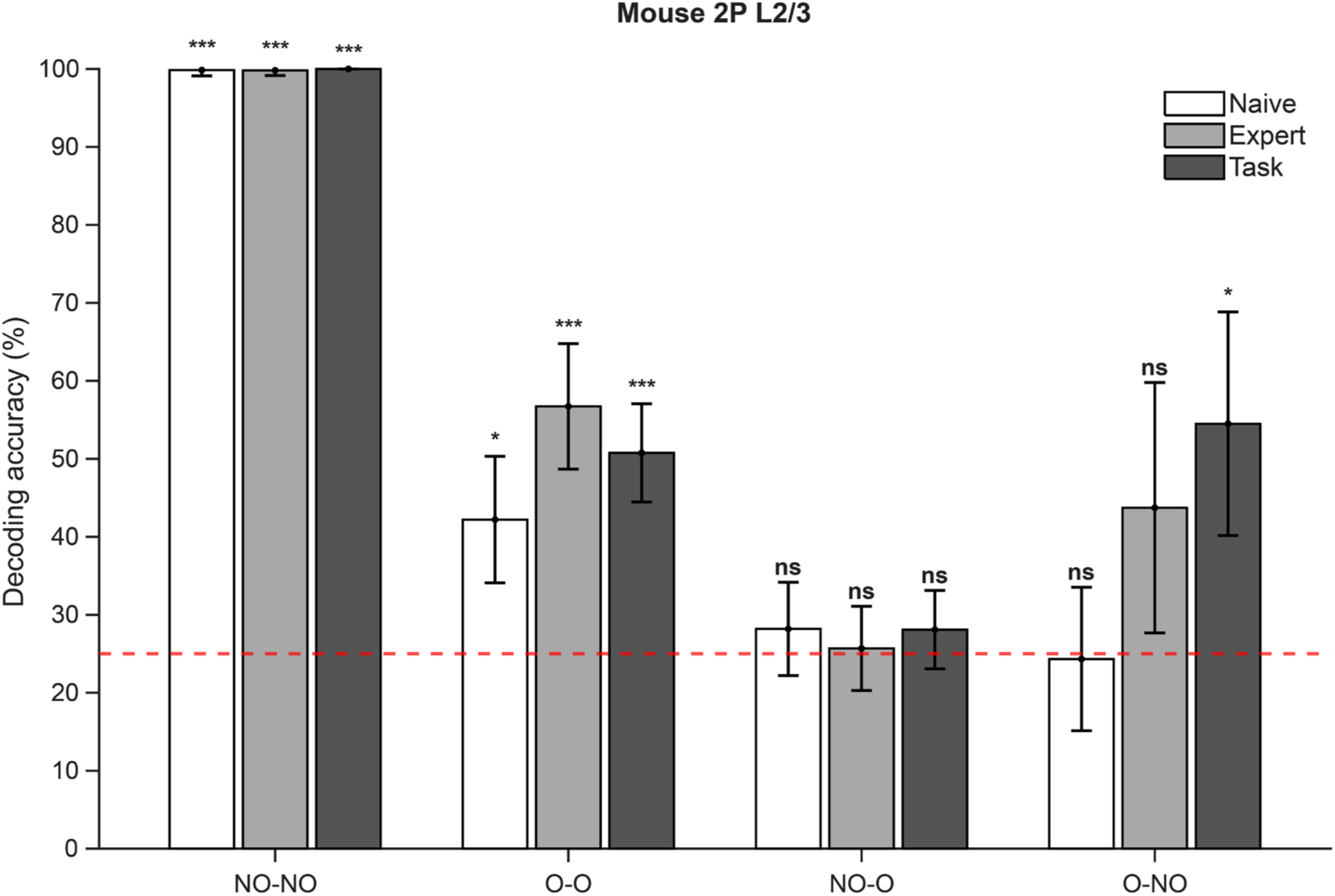
Decoding of visual and contextual signals across conditions. Accuracy for population decoding of stimulus identity for mouse L2/3 neurons across all conditions. Linear discriminant analysis (LDA) decoders were trained and tested on independent trial sets for within-condition decoding (NO-NO, O-O) and cross-condition decoding (NO-O, O-NO). To control for differences in population size across conditions, each decoding run was performed on a randomly selected subset of 300 neurons, corresponding to the smallest available population size across experimental groups. Data were split equally into training and test sets, and decoding performance was averaged across repeated subsampling iterations. Bars show mean ± SD across decoding repetitions. Chance performance (red dashed line) was 25%. Statistical significance was assessed using permutation tests. ***p < 0.001, **p < 0.01, *p < 0.05, ns: not significant. NO: non-occluded; O: occluded.

**Fig. S14.**
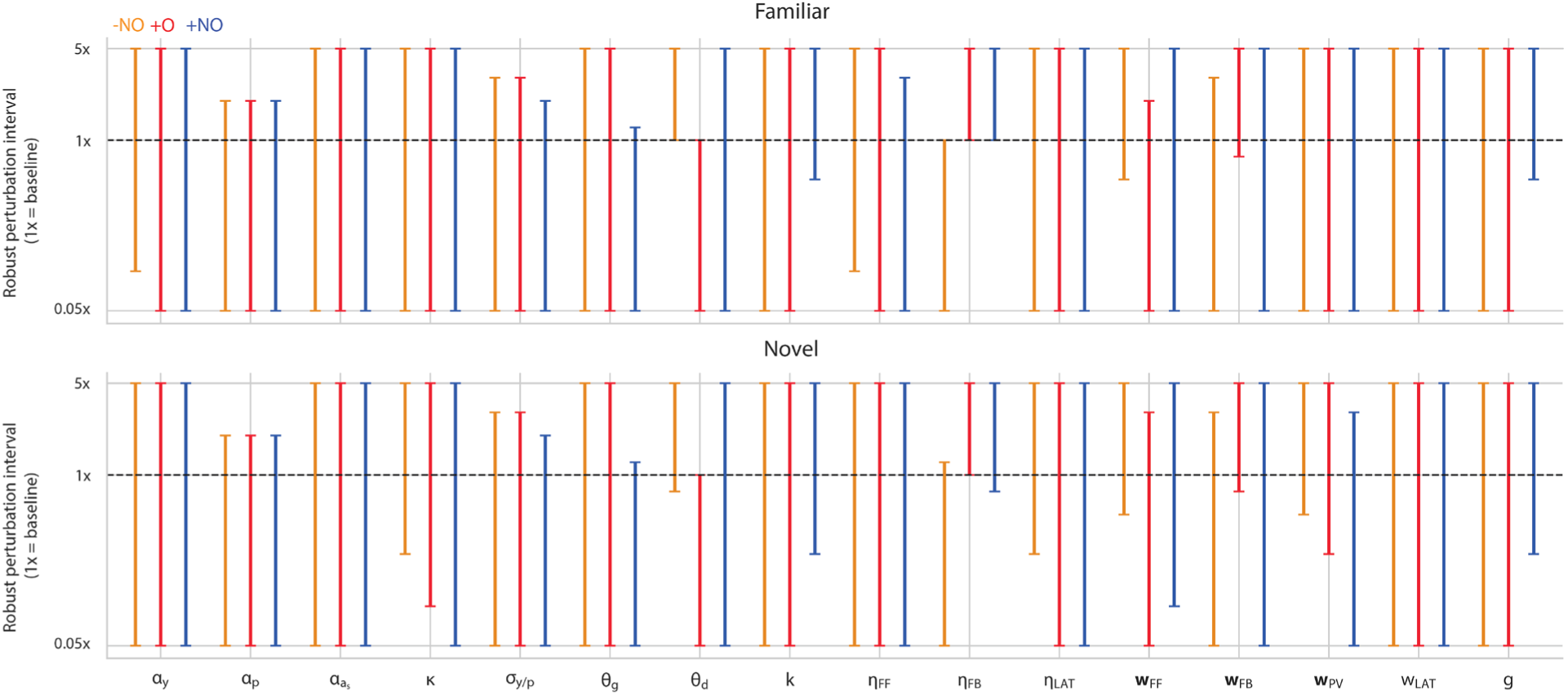
Robustness of computational model to parameter perturbation. Each parameter was perturbed in the range between 0.05x and 5x of its fitted value. The interval of perturbations (each with different random seeds) over which the transition sector of each model’s transition vector remained the same, and retained at least 25% of its magnitude is displayed for each parameter.

**Figure S15:**
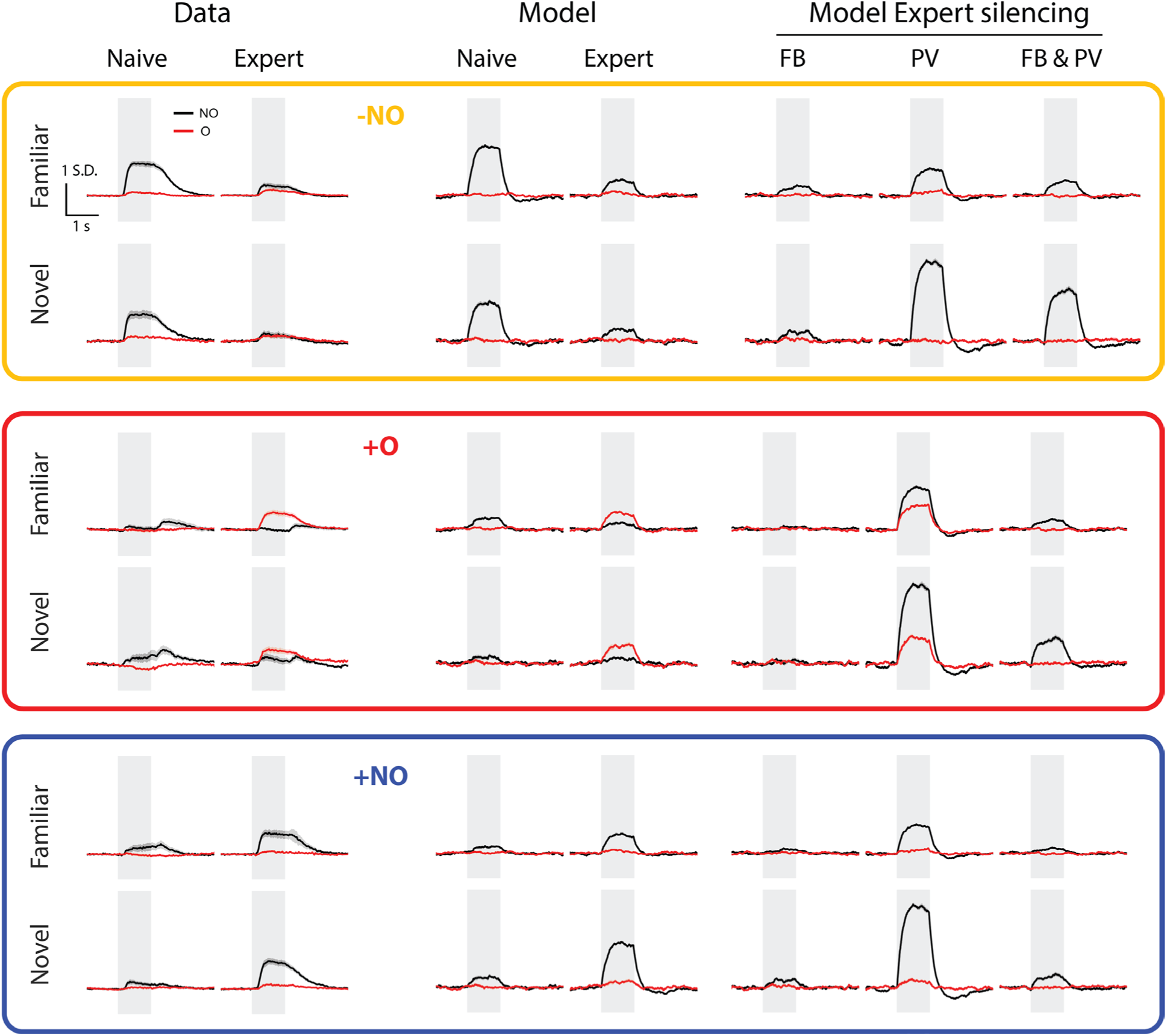
Neural responses for expert ablation experiments in computational model Dominant modes of experience-dependent plasticity under silencing experiments. Response traces indicate average over all recorded neurons undergoing either –NO, +O, or +NO transition. For model these response traces are average over 10 random seeds, for each of the six mean field models (separate parameter set fitted for each transition in familiar and novel).

**Table S1.** Shared parameters.

| Symbol | Value | Meaning |
| --- | --- | --- |
| $\alpha_y$ | 0.05 | PyC leak rate |
| $\alpha_p$ | 0.5 | PV leak rate |
| $\alpha_{af}$ | 0.01 | Fast adaptation rate |
| $\alpha_{as}$ | 0.0025 | Slow FF-activity accumulator rate |
| $\kappa$ | 10 | Divisive normalization scale |
| $\sigma_{y/p}$ | 0.2 | PyC and PV baseline-current s.d. |
| $\theta_g$ | 0.1 | Apical gain threshold |
| $\theta_d$ | 0.14 | Apical drive threshold |
| $k$ | 10 | Apical-gain sigmoid steepness |
| $\eta_{FF}$ | 5.0 | FF depression rate per step |
| $\eta_{FB}$ | 0.0003 | FB potentiation rate per step |
| $\eta_{LAT}$ | 0.05 | PV-to-PyC potentiation rate per step |
| $x_i$ | $\text{perm}_i([1,0,0])$ | Feedforward input |
| $c_i$ | $\text{perm}_i([1,.6,.6])/2.2$ | Contextual input |
| $T$ | 400 | Steps per trial (100 steps/s) |
| $N_{train}$ | 7 | Training trials per familiar image |
| $N_{test}$ | 50 | Naive and expert trials per image |

**Table S2.** Initial parameters defining the six mean-field transition models.

| Source | Transition | $w_{FF}$ | $w_{FB}$ | $w_{PV}$ | $w_{LAT}$ | $g$ |
| --- | --- | --- | --- | --- | --- | --- |
| Familiar | $-NO$ | (.05,.05,.05) | (.0125,.0125,.0125) | (.1,.1,.1) | .025 | 5 |
| Familiar | $+O$ | (.015,.015,.015) | (.035,.035,.035) | (.6,.6,.6) | .05 | 5 |
| Familiar | $+NO$ | (.015,.0015,.0015) | (.0125,.0125,.0125) | (.05,.05,.05) | .0125 | 35 |
| Novel | $-NO$ | (.05,.05,.05) | (.0125,.0125,.0125) | (.5,.5,.5) | .05 | 5 |
| Novel | $+O$ | (.025,.025,.025) | (.045,.045,.045) | (1.0,1.0,1.0) | .2 | 5 |
| Novel | $+NO$ | (.015,.015,.015) | (.025,.025,.025) | (.1,.1,.1) | .025 | 35 |

## REFERENCES

1. Zmarz, P. & Keller, G. B. Mismatch Receptive Fields in Mouse Visual Cortex. Neuron 92, 766–772 (2016).

2. Lamme, V. A. The neurophysiology of figure-ground segregation in primary visual cortex. J Neurosci 15, 1605–1615 (1995).

3. Homann, J., Koay, S. A., Chen, K. S., Tank, D. W. & Berry, M. J. Novel stimuli evoke excess activity in the mouse primary visual cortex. Proc Natl Acad Sci U S A 119, e2108882119 (2022).

4. Furutachi, S., Franklin, A. D., Aldea, A. M., Mrsic-Flogel, T. D. & Hofer, S. B. Cooperative thalamocortical circuit mechanism for sensory prediction errors. Nature 633, 398–406 (2024).

5. Hamm, J. P. & Yuste, R. Somatostatin Interneurons Control a Key Component of Mismatch Negativity in Mouse Visual Cortex. Cell Reports 16, 597–604 (2016).

6. Mendoza-Halliday, D., Xu, H., Azevedo, F. A. C. & Desimone, R. Dissociable neuronal substrates of visual feature attention and working memory. Neuron 112, 850–863.e6 (2024).

7. Keller, G. B., Bonhoeffer, T. & Hübener, M. Sensorimotor mismatch signals in primary visual cortex of the behaving mouse. Neuron 74, 809–815 (2012).

8. Clark, A. Whatever next? Predictive brains, situated agents, and the future of cognitive science. Behav Brain Sci 36, 181–204 (2013).

9. Rao, R. P. & Ballard, D. H. Predictive coding in the visual cortex: a functional interpretation of some extra-classical receptive-field effects. Nat Neurosci 2, 79–87 (1999).

10. Friston, K. Functional integration and inference in the brain. Prog Neurobiol 68, 113–143 (2002).

11. Keller, G. B. & Mrsic-Flogel, T. D. Predictive Processing: A Canonical Cortical Computation. Neuron 100, 424–435 (2018).

12. Bastos, A. M. et al. Canonical microcircuits for predictive coding. Neuron 76, 695–711 (2012).

13. Shipp, S. Neural Elements for Predictive Coding. Front Psychol 7, 1792 (2016).

14. Jordan, R. & Keller, G. B. Opposing Influence of Top-down and Bottom-up Input on Excitatory Layer 2/3 Neurons in Mouse Primary Visual Cortex. Neuron 108, 1194–1206.e5 (2020).

15. Liu, Y. et al. Organization of corticocortical and thalamocortical top-down inputs in the primary visual cortex. Nat Commun 15, 4495 (2024).

16. Larkum, M. A cellular mechanism for cortical associations: an organizing principle for the cerebral cortex. Trends in Neurosciences 36, 141–151 (2013).

17. Allman, J., Miezin, F. & McGuinness, E. Stimulus Specific Responses from Beyond the Classical Receptive Field: Neurophysiological Mechanisms for Local-Global Comparisons in Visual Neurons. Annual Review of Neuroscience 8, 407–430 (1985).

18. Niell, C. M. & Stryker, M. P. Modulation of Visual Responses by Behavioral State in Mouse Visual Cortex. Neuron 65, 472–479 (2010).

19. Iurilli, G. et al. Sound-Driven Synaptic Inhibition in Primary Visual Cortex. Neuron 73, 814–828 (2012).

20. Meijer, G. T., Montijn, J. S., Pennartz, C. M. A. & Lansink, C. S. Audiovisual Modulation in Mouse Primary Visual Cortex Depends on Cross-Modal Stimulus Configuration and Congruency. J Neurosci 37, 8783–8796 (2017).

21. Muckli, L. & Petro, L. S. Network interactions: non-geniculate input to V1. Current Opinion in Neurobiology 23, 195–201 (2013).

22. Muckli, L. et al. Contextual Feedback to Superficial Layers of V1. Current Biology 25, 2690–2695 (2015).

23. Morgan, A. T., Petro, L. S. & Muckli, L. Scene Representations Conveyed by Cortical Feedback to Early Visual Cortex Can Be Described by Line Drawings. J. Neurosci. 39, 9410–9423 (2019).

24. Revina, Y., Petro, L. S. & Muckli, L. Cortical feedback signals generalise across different spatial frequencies of feedforward inputs. NeuroImage 180, 280–290 (2018).

25. Smith, F. W. & Muckli, L. Nonstimulated early visual areas carry information about surrounding context. Proc. Natl. Acad. Sci. U.S.A. 107, 20099–20103 (2010).

26. Petro, L. S., Smith, F. W., Abbatecola, C. & Muckli, L. The Spatial Precision of Contextual Feedback Signals in Human V1. Biology 12, 1022 (2023).

27. Papale, P. et al. The representation of occluded image regions in area V1 of monkeys and humans. Current Biology 33, 3865–3871.e3 (2023).

28. Sawtell, N. B. et al. NMDA Receptor-Dependent Ocular Dominance Plasticity in Adult Visual Cortex. Neuron 38, 977–985 (2003).

29. Dumoulin, S. O. & Wandell, B. A. Population receptive field estimates in human visual cortex. NeuroImage 39, 647–660 (2008).

30. Erisken, S. et al. Effects of Locomotion Extend throughout the Mouse Early Visual System. Current Biology 24, 2899–2907 (2014).

31. Keller, A. J., Roth, M. M. & Scanziani, M. Feedback generates a second receptive field in neurons of the visual cortex. Nature 582, 545–549 (2020).

32. Kirchberger, L., Mukherjee, S., Self, M. W. & Roelfsema, P. R. Contextual drive of neuronal responses in mouse V1 in the absence of feedforward input. Sci Adv 9, eadd2498 (2023).

33. Makino, H. & Komiyama, T. Learning enhances the relative impact of top-down processing in the visual cortex. Nat Neurosci 18, 1116–1122 (2015).

34. Muzzu, T. & Saleem, A. B. Feature selectivity can explain mismatch signals in mouse visual cortex. Cell Rep 38, 110413 (2022).

35. Phillips, W. A. et al. Cellular psychology: relating cognition to context-sensitive pyramidal cells. Trends in Cognitive Sciences 29, 28–40 (2025).

36. Abbott, L. F., Varela, J. A., Sen, K. & Nelson, S. B. Synaptic depression and cortical gain control. Science 275, 220–224 (1997).

37. Tsodyks, M. V. & Markram, H. The neural code between neocortical pyramidal neurons depends on neurotransmitter release probability. Proc Natl Acad Sci U S A 94, 719–723 (1997).

38. Grubb, M. S. & Burrone, J. Activity-dependent relocation of the axon initial segment fine-tunes neuronal excitability. Nature 465, 1070–1074 (2010).

39. Evans, M. D., Dumitrescu, A. S., Kruijssen, D. L. H., Taylor, S. E. & Grubb, M. S. Rapid Modulation of Axon Initial Segment Length Influences Repetitive Spike Firing. Cell Rep 13, 1233–1245 (2015).

40. Desai, N. S., Rutherford, L. C. & Turrigiano, G. G. Plasticity in the intrinsic excitability of cortical pyramidal neurons. Nat Neurosci 2, 515–520 (1999).

41. Ibata, K., Sun, Q. & Turrigiano, G. G. Rapid synaptic scaling induced by changes in postsynaptic firing. Neuron 57, 819–826 (2008).

42. Xue, M., Atallah, B. V. & Scanziani, M. Equalizing excitation–inhibition ratios across visual cortical neurons. Nature 511, 596–600 (2014).

43. Schulz, A., Miehl, C., Berry, M. J. & Gjorgjieva, J. The generation of cortical novelty responses through inhibitory plasticity. Elife 10, e65309 (2021).

44. Kok, P., Jehee, J. F. M. & de Lange, F. P. Less Is More: Expectation Sharpens Representations in the Primary Visual Cortex. Neuron 75, 265–270 (2012).

45. Freedman, D. J., Riesenhuber, M., Poggio, T. & Miller, E. K. Experience-dependent sharpening of visual shape selectivity in inferior temporal cortex. Cereb Cortex 16, 1631–1644 (2006).

46. Grill-Spector, K., Henson, R. & Martin, A. Repetition and the brain: neural models of stimulus-specific effects. Trends in Cognitive Sciences 10, 14–23 (2006).

47. Carandini, M. & Heeger, D. J. Normalization as a canonical neural computation. Nat Rev Neurosci 13, 51–62 (2011).

48. Kumar, S., Kaposvari, P. & Vogels, R. Encoding of Predictable and Unpredictable Stimuli by Inferior Temporal Cortical Neurons. Journal of Cognitive Neuroscience 29, 1445–1454 (2017).

49. De Lange, F. P., Heilbron, M. & Kok, P. How Do Expectations Shape Perception? Trends in Cognitive Sciences 22, 764–779 (2018).

50. Richter, D., Heilbron, M. & De Lange, F. P. Dampened sensory representations for expected input across the ventral visual stream. Oxford Open Neuroscience 1, kvac013 (2022).

51. Blank, H. & Davis, M. H. Prediction Errors but Not Sharpened Signals Simulate Multivoxel fMRI Patterns during Speech Perception. PLoS Biol 14, e1002577 (2016).

52. Han, B., Mostert, P. & De Lange, F. P. Predictable tones elicit stimulus-specific suppression of evoked activity in auditory cortex. NeuroImage 200, 242–249 (2019).

53. Wright, W. J., Hedrick, N. G. & Komiyama, T. Distinct synaptic plasticity rules operate across dendritic compartments in vivo during learning. Science 388, 322–328 (2025).

54. Gambino, F. et al. Sensory-evoked LTP driven by dendritic plateau potentials in vivo. Nature 515, 116–119 (2014).

55. Bastos, G. et al. Top-down input modulates visual context processing through an interneuron-specific circuit. Cell Rep 42, 113133 (2023).

56. Williams, L. E. & Holtmaat, A. Higher-Order Thalamocortical Inputs Gate Synaptic Long-Term Potentiation via Disinhibition. Neuron 101, 91–102.e4 (2019).

57. van Versendaal, D. & Levelt, C. N. Inhibitory interneurons in visual cortical plasticity. Cellular and Molecular Life Sciences 73, 3677–3691 (2016).

58. Cuevas, N. et al. Stimulus-specificity of surround-induced responses in primary visual cortex. eLife 13, (2025).

59. Hofer, S. B. et al. Differential connectivity and response dynamics of excitatory and inhibitory neurons in visual cortex. Nat Neurosci 14, 1045–1052 (2011).

60. Shin, H. et al. Recurrent pattern completion drives the neocortical representation of sensory inference. Nat Neurosci 28, 2319–2329 (2025).

61. Major, G., Larkum, M. E. & Schiller, J. Active properties of neocortical pyramidal neuron dendrites. Annu Rev Neurosci 36, 1–24 (2013).

62. Larkum, M. E., Zhu, J. J. & Sakmann, B. A new cellular mechanism for coupling inputs arriving at different cortical layers. Nature 398, 338–341 (1999).

63. Larkum, M. E., Waters, J., Sakmann, B. & Helmchen, F. Dendritic spikes in apical dendrites of neocortical layer 2/3 pyramidal neurons. J Neurosci 27, 8999–9008 (2007).

64. Marvan, T. & Phillips, W. A. Cellular mechanisms of cooperative context-sensitive predictive inference. Curr Res Neurobiol 6, 100129 (2024).

65. Marvan, T., Polák, M., Bachmann, T. & Phillips, W. A. Apical amplification—a cellular mechanism of conscious perception? Neurosci Conscious 2021, niab036 (2021).

66. Dias, R. F. et al. Visual experience reduces the spatial redundancy between cortical feedback inputs and primary visual cortex neurons. Neuron 112, 3329–3342.e7 (2024).

67. Rajan, R. et al. Visual experience exerts an instructive role on cortical feedback inputs to the primary visual cortex. Curr Biol S0960-9822(26)00070–9 (2026) doi:10.1016/j.cub.2026.01.031.

68. Summerfield, C., Wyart, V., Johnen, V. M. & De Gardelle, V. Human Scalp Electroencephalography Reveals that Repetition Suppression Varies with Expectation. Front. Hum. Neurosci. 5, (2011).

69. Meyer, T. & Olson, C. R. Statistical learning of visual transitions in monkey inferotemporal cortex. Proc. Natl. Acad. Sci. U.S.A. 108, 19401–19406 (2011).

70. Todorovic, A. & De Lange, F. P. Repetition Suppression and Expectation Suppression Are Dissociable in Time in Early Auditory Evoked Fields. J. Neurosci. 32, 13389–13395 (2012).

71. Wacongne, C., et al. Evidence for a hierarchy of predictions and prediction errors in human cortex. Proc. Natl. Acad. Sci. U.S.A. 108, 20754–20759 (2011).

72. Von Helmholtz, H. Handbuch der physiologischen Optik. (1867).

73. Keller, G. B. & Sterzer, P. Predictive Processing: A Circuit Approach to Psychosis. Annu Rev Neurosci 47, 85–101 (2024).

74. Hebart, M. N. et al. THINGS: A database of 1,854 object concepts and more than 26,000 naturalistic object images. PLOS ONE 14, e0223792 (2019).

75. Van Beest, E. H. et al. Mouse visual cortex contains a region of enhanced spatial resolution. Nat Commun 12, 4029 (2021).

76. De Kraker, L. et al. SpecSeg is a versatile toolbox that segments neurons and neurites in chronic calcium imaging datasets based on low-frequency cross-spectral power. Cell Reports Methods 2, 100299 (2022).

77. Pnevmatikakis, E. A. & Giovannucci, A. NoRMCorre: An online algorithm for piecewise rigid motion correction of calcium imaging data. Journal of Neuroscience Methods 291, 83–94 (2017).

78. Poort, J. et al. The Role of Attention in Figure-Ground Segregation in Areas V1 and V4 of the Visual Cortex. Neuron 75, 143–156 (2012).

79. Walt, S. van der et al. scikit-image: image processing in Python. PeerJ 2, e453 (2014).

80. Knijnenburg, T. A., Wessels, L. F. A., Reinders, M. J. T. & Shmulevich, I. Fewer permutations, more accurate P-values. Bioinformatics 25, i161–168 (2009).

81. Larkum, M. E., Senn, W. & Lüscher, H.-R. Top-down dendritic input increases the gain of layer 5 pyramidal neurons. Cereb Cortex 14, 1059–1070 (2004).

82. Syeda, A. et al. Facemap: a framework for modeling neural activity based on orofacial tracking. Nat Neurosci 27, 187–195 (2024).

